# The Molecular Genetic Basis of Herbivory between Butterflies and their Host-Plants

**DOI:** 10.1101/154799

**Authors:** Sumitha Nallu, Jason Hill, Kristine Don, Carlos Sahagun, Wei Zhang, Camille Meslin, Emilie Snell-Rood, Nathan L. Clark, Nathan I. Morehouse, Joy Bergelson, Christopher W. Wheat, Marcus R. Kronforst

## Abstract

Interactions between herbivorous insects and their host-plants are a central component of terrestrial food webs and a critical topic in agriculture, where a substantial fraction of potential crop yield is lost annually to pests. Important insights into plant-insect interactions have come from research on specific plant defenses and insect detoxification mechanisms. Yet, much remains unknown about the molecular mechanisms that mediate plant-insect interactions. Here we use multiple genome-wide approaches to map the molecular basis of herbivory from both plant and insect perspectives, focusing on butterflies and their larval host-plants. Parallel genome-wide association studies in the Cabbage White butterfly, *Pieris rapae*, and its host-plant, *Arabidopsis thaliana*, pinpointed a small number of butterfly and plant genes that influenced herbivory. These genes, along with much of the genome, were regulated in a dynamic way over the time course of the feeding interaction. Comparative analyses, including diverse butterfly/plant systems, showed a variety of genome-wide responses to herbivory, yet a core set of highly conserved genes in butterflies as well as their host-plants. These results greatly expand our understanding of the genomic causes and evolutionary consequences of ecological interactions across two of Nature’s most diverse taxa, butterflies and flowering plants.

## Introduction

Butterflies and moths, and the host-plants that their larvae feed on, comprise two of the largest groups of species on earth, the Lepidoptera and Angiosperms. The extreme diversity of these two groups has arisen from the co-evolutionary interactions between them, wherein evolution proceeded via reciprocal adaptations as each clade evolved in response to changes in the other^1,2^. In their seminal paper, Ehrlich and Raven^3^ used the diffuse evolutionary relationships among butterflies and their host-plants to formally introduce the concept of co-evolution. In the 50 years since Ehrlich and Raven^3^, research on the molecular basis of herbivory between butterflies and their host-plants has revealed important insights into specific chemicals and defense pathways utilized by plants, such as furanocoumarins, glucosinolates, and cardenolides^4^. Research on butterflies, in turn, has revealed specific genes and gene families involved in host-plant detoxification, such as cytochrome P450 enzymes^5,6^, the nitrile specifier protein of pierid butterflies^7^, and the Na(+), K(+) –ATPase of the monarch butterfly^8^. Yet, despite these important advances, our understanding of the molecular genetic basis of these co-evolutionary interactions remains limited. Genomic approaches, including genome-wide association (GWA) studies and transcriptomics, provide a means to move beyond candidate genes and pathways to uncover the molecular determinants of this fundamental ecological interaction in an unbiased way^9,10^. Here we utilize these methods in a diversity of butterflies and their respective host-plant to uncover the genetic basis of herbivory.

## Results

### Genome wide associations with herbivory: the host-plant, *Arabidopsis thaliana*

First, we investigated the genetics of butterfly/host herbivory by mapping associated variants in parallel GWA studies focused on the flowering plant *Arabidopisis thaliana* and its natural insect herbivore, the Cabbage White butterfly, *Pieris rapae*. For these experiments, we used either 96 natural accessions of *A. thaliana* and a single lab strain of *P. rapae* (for the plant GWAS), or the offspring of 96 field-caught females of *P. rapae* and a single accession of *A. thaliana* (for the butterfly GWAS). For both experiments, we measured herbivory as the amount of weight gained and the amount of leaf surface area eaten by second instar larvae over a period of 72 hours. *Arabidopisis thaliana* GWA resulted in a total of 90 associated SNPs that were in linkage-disequilibrium with 389 genes, a gene set enriched for plant defense (Supplementary data 1). A subset of 12 well-supported candidate genes contained three or more associated SNPs each (Fig. 1a), eight of which we were able to functionally validate using SALK T-DNA mutants (Fig. 1c). This validated gene set includes well-known and novel defense genes. For instance, the cytochrome P450 gene CYP79B2 is involved in the conversion of tryptophan to indole-3- acetaldoxime, a precursor of indole glucosinolates and indole-3-acetic acid^11^. Indole glucosinolates are important secondary metabolites used for defense by *Arabidopsis*, and other species in the plant family Brassicaceae, and they have been shown to deter herbivores and pathogens ^12,13^. Insects that feed on Brassicaceae have evolved various physiological strategies against toxic effects of glucosinolates^14^. The larvae of *P. rapae*, for instance, redirect the hydrolysis pathway catalyzed by myrosinase^7^, and instead of producing toxic isothiocyanates, hydrolysis is redirected toward the formation of nitriles by the butterfly’s nitrile specifier protein (NSP)^15,16^. Another functionally validated gene, phosphoribosylanthranilate isomerase 3 (PAI3), catalyzes a step in the L-tryptophan synthesis pathway^17^, to produce the precursor of indole glucosionolates, although there is no direct evidence that it influences glucosinolates. The genes PROPEP1 and PROPEP3 belong to the AtPep (endogenous danger peptides) gene family. They are associated with activation of danger- or damage-associated molecular patterns (DAMPs) immunity in plants against both pathogen and herbivore attack^18^. The other four genes validated in our study do not have known roles in defense and include an uncharacterized cytochrome P450 (CYP705A33), Importin alpha (IMPA-1), a CTP synthase (AT4G20320), and an S-adenosyl-L-methionine-dependent methyltransferase (AT1G29470). Interestingly, the *A. thaliana* genes we found associated with herbivory did not overlap with those from a recent genome-wide association study of methionine-derived glucosinolates in *A. thaliana*^19^.

**Figure 1:**
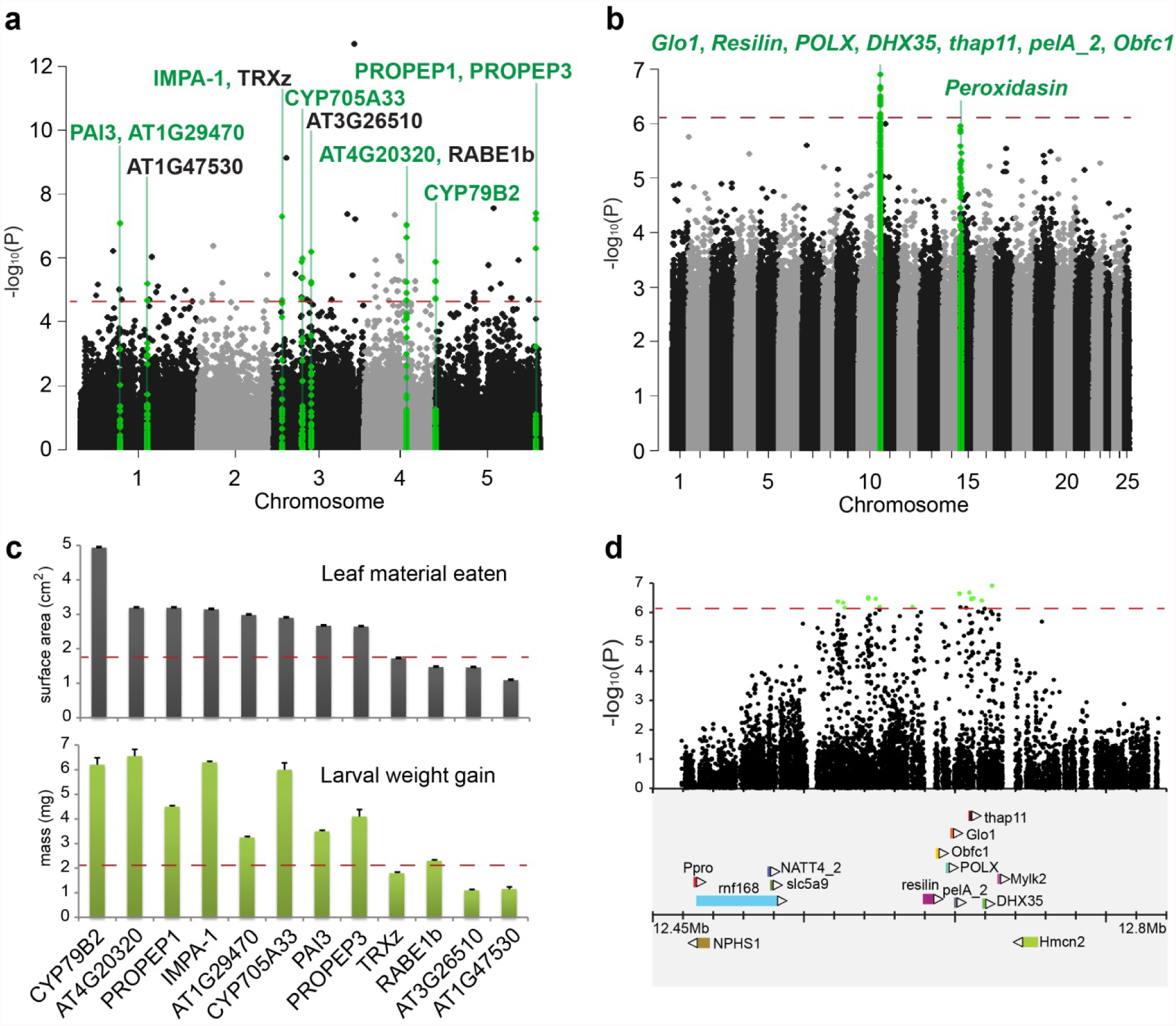
Butterfly and host-plant herbivory candidate genes identified by parallel genomewide association studies. (a) Manhattan plot with the associated SNPs and their corresponding P-values from the herbivory GWAS based on 96 accessions of *Arabidopsis thaliana*. The eight genes that were functionally validated among the primary 12 candidate genes are highlighted in green. (b) Manhattan plot with the associated SNPs and their corresponding P-values from the herbivory GWAS based on 96 *Pieris rapae* samples. The genes contained in the two association peaks are listed. The significance threshold for each GWAS is indicated with a dashed red line. (c) Average leaf surface area eaten (top panel) and weight gained (bottom panel) by larvae on the SALK T-DNA mutants for the 12 candidate genes from the *Arabdiopsis* GWAS. The corresponding value from the wild type plant (col-0) is represented as a line across each histogram. (d) Zoom-in on the chromosome 10 region associated with herbivory in the *P. rapae* genome. Significantly associated SNPs are indicated in green with a gene map below.

### *Pieris rapae* genome sequencing

We began our butterfly GWAS by assembling a high-quality reference genome sequence for the Cabbage White butterfly, *Pieris rapae*. For the reference genome, we combined next-generation DNA sequencing data from one PCR-free paired-end Illumina library, three mate-pair libraries (3 kbp, 7 kbp, 40 kbp), and a Dovetail Chicago library to generate a final assembly of 320 Mbp, with a N50 of 11.5 Mbp, spanning the 25 *P. rapae* chromosomes^20^. Annotation was performed using a diverse set of RNA-Seq data. This chromosomal-level assembly of the butterfly genome (Supplementary Fig. 1) provided an essential genomic foundation with which to explore the insect side of the plant-insect interaction.

### Genome wide associations with herbivory: the herbivore, *Pieris rapae*

For GWAS, we performed our larval feeding trials with the offspring of wild-caught females and used whole genome resequencing to genotype 96 unrelated larvae at genome-wide single-nucleotide polymorphisms (SNPs). *Pieris rapae* GWA revealed just two strongly associated regions in the genome (Fig. 1b), the largest of which encompassed a total of 16 significantly associated SNPs distributed across 98 kbp on chromosome 10, spanning seven genes (Fig. 1d). One of these genes, *Glyoxalase 1* (*Glo1*), a lactoylglutathione lyase, is a central enzyme in the glyoxalase pathway present in all organisms. Glyoxalase detoxifies cytosolic methyglyoxyl, a toxic by-product of metabolism, in a two-step process that utilizes glutathione. Glutathione itself is an important metabolite involved in multiple biological processes in plants and animals^21^. In addition to *Glo1*, the 98 kbp associated region also contained the following genes: *Resilin*, a Retrovirus-related Pol polyprotein POLX, the ATP-dependent RNA helicase *DHX35*, a THAP domain-containing protein, and two uncharacterized genes. A second associated region in the genome, on chromosome 15, contained just one gene, *peroxidasin* (Supplementary Fig. 2). Peroxidasin genes play important roles in morphogenesis, cell adhesion, formation of extracellular matrix and defense against pathogens^22,23^. Our results suggest this gene plays a role in larval growth, possibly in the context of host-plant detoxification.

### Herbivory-induced differential genes expression experiments

To further interrogate the GWA genes, and explore genome-wide patterns of gene expression throughout the plant-insect interaction, we next used transcriptomics to measure differential gene expression in *P. rapae* and *A. thaliana* over the time course of their interaction. For these experiments, butterfly and plant were allowed to interact for a period of time, after which we harvested tissue from both organisms, as well as controls, for RNA-seq. For these experiments, the plant and butterfly genome sequences were used as references to analyze RNA-seq data. To capture gene expression changes in response to oviposition, we compared gene expression profiles of (a) leaves with eggs (Fig. 2a,b) vs. leaves without eggs 72 h after oviposition, and (b) eggs deposited on leaves vs. eggs deposited on wax paper 72 h after oviposition (Fig. 2c). To capture gene expression changes of the plant in response to larval feeding, we compared (c) leaves after 24 h of larval feeding from plants previously exposed to eggs vs. leaves after 24 h of larval feeding from plants not previously exposed to eggs (Fig. 2d) vs. control leaves never exposed to eggs or larvae. We also analyzed leaves 24 h after mechanical wounding (Fig. 2e) to compare gene expression changes associated with larval feeding vs. wounding. To identify gene expression changes in the insect associated with feeding, we compared (d) larvae after 24 h of feeding on leaves from plants exposed to eggs vs. larvae after 24 h of feeding on leaves from plants not exposed to eggs vs. larvae after 24 h of feeding on artificial diet (Fig. 2f). We also measured gene expression in butterfly pupae and adults, for comparison.

**Figure 2:**
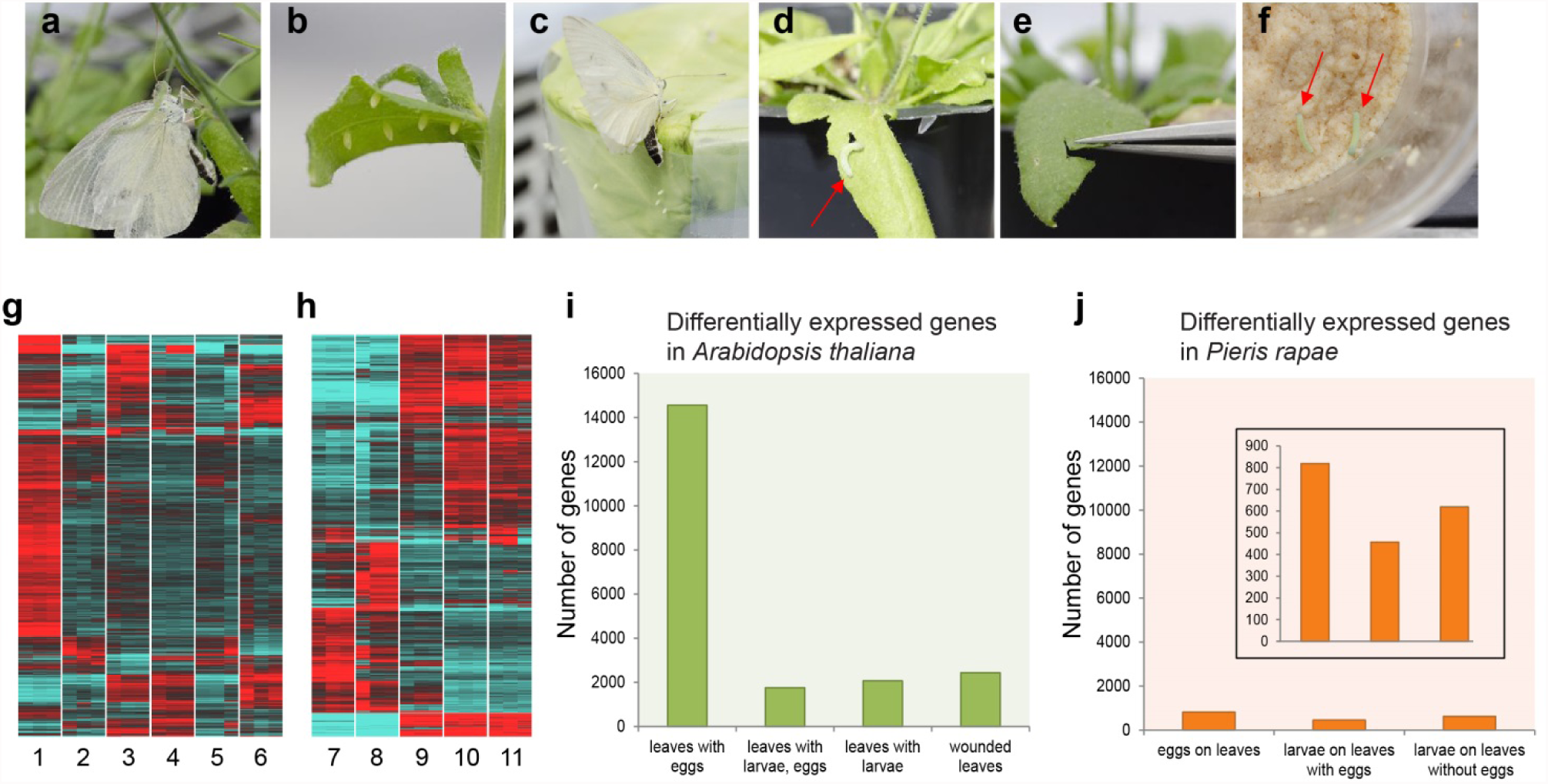
Genome-wide responses to herbivory in *Arabidopsis thaliana* and *Pieris rapae* over the time course of their interaction. Samples collected from various stages of interaction: (a, b) eggs laid on the leaves, (c) eggs laid on the wax paper, (d) larva feeding on a leaf, (e) mechanical wounding of the leaf, (f) larvae feeding on artificial diet. Heat maps of differentially expressed genes in (g) *Arabidopsis thaliana* and (h) *Pieris rapae*. Color scale ranges from ≤ -1.5 log fold change (blue) to ≥1.5 log fold change (red). Individual treatments correspond to: 1) leaves with eggs 72h after oviposition, 2) leaves with no eggs, control for 72h oviposition treatment, 3) leaves after 24h larval (48h old) feeding, 4) leaves with eggs after 24h larval (48h old) feeding, 5) leaves with no eggs and larvae, control for larval feeding and wounding, 6) leaves 24h after wounding, 7) eggs 72h after oviposition on wax paper, 8) eggs 72h after oviposition on leaves, 9) 48h old larvae after 24h feeding on artificial diet, 10) 48h old larvae after 24h feeding on leaves with eggs, 11) 48h old larvae after 24h feeding on leaves. (i) Number of differentially expressed *Arabidopsis thaliana* genes across treatments. (j) Number of differentially expressed *Pieris rapae* genes across treatments - the inset shows the same results plotted with a reduced y-axis scale.

The genes identified in our butterfly and plant GWA experiments showed patterns of expression consistent with a role in herbivory (Fig. 3). In *A. thaliana*, eight of 12 genes identified in the GWA study had the highest expression in leaves exposed to eggs and larvae, including both cytochrome P450 genes and PROPEP3. PROPEP1 and PAI3 showed elevated expression in leaves with eggs only, followed by leaves with larvae only. The remaining two genes, IMPA-1 and RABE1b (one of the four genes that was not validated) had other expression patterns, showing elevated expression in mechanically wounded leaves and the leaf control, respectively (Fig. 3a). In *P. rapae*, six of the seven genes in the associated region on chromosome 10, and *peroxidasin* on chromosome 15 (Supplementary Fig. 2), had the highest expression in larval treatments—the larva is the herbivorous life stage of the butterfly—and were expressed at lower levels in eggs, pupae and adults. Notably, this pattern of expression was distinct from physically adjacent genes that were located outside the associated regions (Fig. 3b and 3c). The one exception, POLX, which was not a candidate gene based on function, showed similar levels of expression across developmental stages. *Resilin* expression was elevated specifically in larvae feeding on leaves, as opposed to those eating artificial diet, indicative of a role directly in the plant-insect interaction.

**Figure 3:**
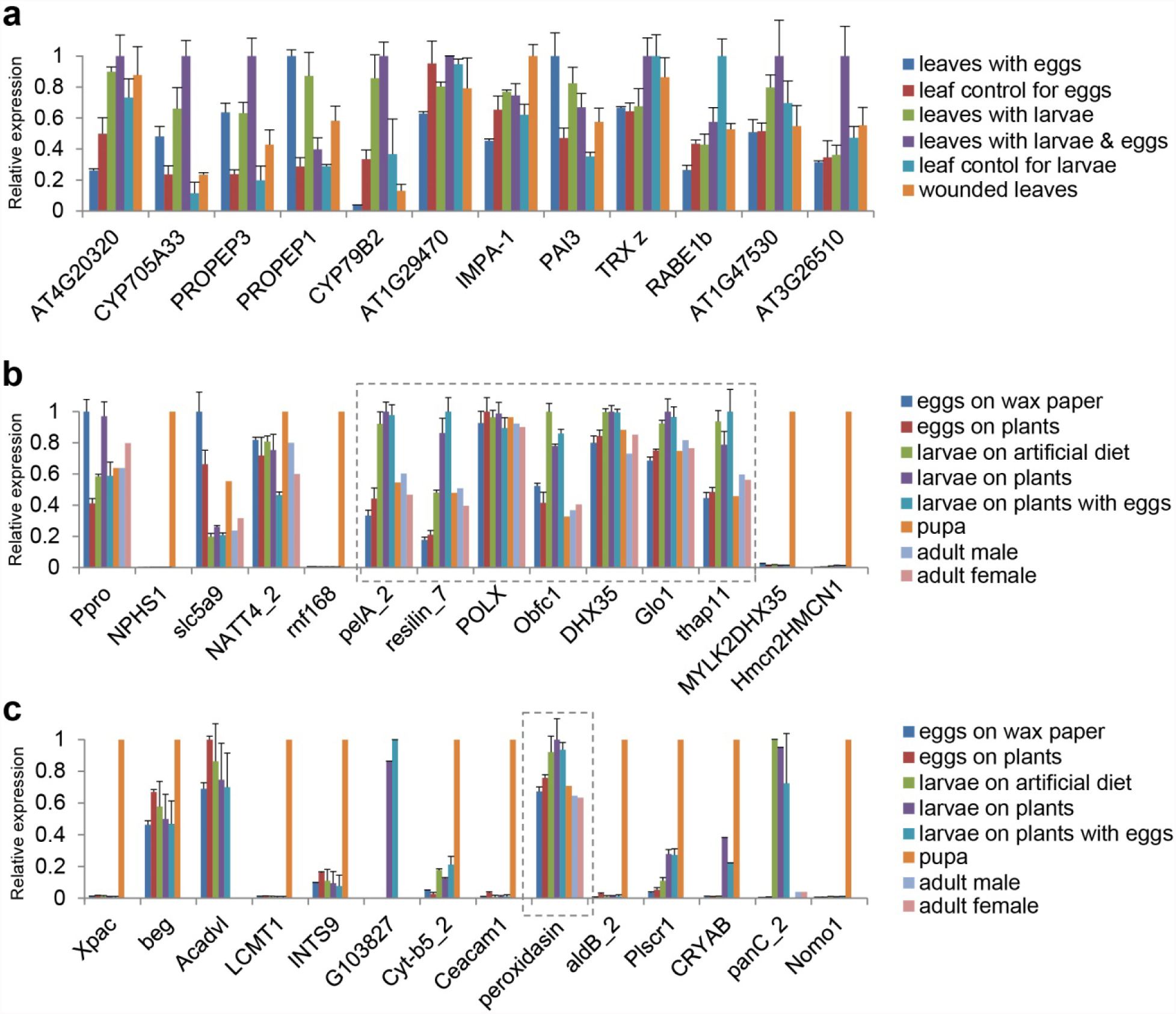
Temporal expression patterns of herbivory candidate genes in *Arabidopsis thaliana* and *Pieris rapae*. . (a) Expression patterns from various RNA-seq treatments of the 12 *Arabidopsis thaliana* candidate genes identified in the herbivory GWAS. (b) Expression patterns from various RNA-seq treatments of the seven *Pieris rapae* genes contained in the associated region on chromosome 10 (boxed), identified in the *Pieris rapae* herbivory GWAS, as well as genes from the adjoining 100kb regions upstream and downstream. (c) Expression patterns from the various RNA-seq treatments of the *peroxidasin* gene identified in the *Pieris rapae* herbivory GWAS, as well as genes from the adjoining 100kb regions upstream and downstream.

The genome-wide responses of *Arabidopisis* and *Pieris*, in terms of differential gene expression, were also striking. In particular, *A. thaliana* showed a massive response to butterfly oviposition with approximately 50% of the plant’s genes changing expression, mostly up-regulated, in response to a female butterfly laying eggs (Fig. 2g). The response of the plant to larval feeding, in contrast, was more modest. The egg is the first life stage of the herbivore that is in contact with the plant and previous studies have shown that plants launch various defenses against insects even before larval hatching^24^. Egg-induced plant defense strategies include plant-mediated desiccation of eggs, egg dropping, egg crushing, and egg killing^24^. We found that a total of 14,563 genes were differentially expressed in *Arabidopsis* leaves after *P. rapae* egg deposition (Fig. 2i). Genes belonging to defense and stress responses were enriched in all treatments (Supplementary data 2). In addition to genes involved in production of glucosinolates and glutathiones, these include protein kinases, proteolytic enzymes, oxidoreductases, peroxidases, NBS-LRR and defense signaling transcription factors that are known to be involved in release of reactive oxygen species, production of pathogenicity-related proteins, activation of systemic acquired resistance, cell wall modification, and programmed cell death in pathogens^25^. These results mirror, and greatly extend, previous work with microarrays and a different species of *Pieris*, *P. brassicae* showing an elevated response of *A. thaliana* to insect oviposition with induction of genes involved in the hypersensitive-like response and pathogenesis-related genes as well as callose and reactive oxygen species accumulation ^26^^-^^28^.

Similar to the plant, *P. rapae* eggs exhibited an elevated response in comparison to the larval stages (Fig. 2h). Of all the genes differentially expressed between eggs oviposited on leaves and wax paper, approximately 50% of them were uncharacterized. Among the genes with predicted functions, genes responding to stress, oxidoreductases, and proteolytic enzymes were abundant (Supplementary data 3). In the next stage of their interaction, larvae feeding on the plant, we analyzed gene expression patterns in larvae feeding on plants exposed to eggs as well as those feeding on plants not previously exposed to eggs. We found that larvae feeding on plants exposed to eggs had a smaller number of differentially expressed genes compared to larvae feeding on plants that were not exposed to eggs (Fig. 2j). This mirrored gene expression patterns in the plant (Fig. 2i), in which the added effect of herbivory after oviposition was less than the effect of herbivory alone. There were 120 differentially expressed larval genes in common between the two treatments, and among the genes with putative functions, genes involved in defense, stress response and proteolysis were overrepresented (Supplementary data 4). Interestingly, glutathione s-transferases (GSTs) frequently show elevated expression in the larvae of Lepidoptera^29^^-^^31^ and other insects^32^, but we did not detect differential expression of GSTs in *P. rapae*.

### Comparative transcriptomics across butterflies and plants

The strong response of *A. thaliana* to butterfly eggs was a particularly striking result. To determine if this was a general property of butterfly-plant interactions, we expanded our analysis of gene expression to three additional plant/insect systems: *Medicago sativa*/*Colias eurytheme*, *Citrofortunella microcarpa*/*Papilio polytes* and *Passiflora oerstedii*/*Heliconius cydno* (Fig. 4a). For this comparative analysis, we generated *de novo* transcriptomes for all organisms to use as references to analyze patterns of differential gene expression using RNA-seq data. We also reanalyzed our *A. thaliana*/*P. rapae* data using *de novo* reference transcriptomes, as opposed to the genome sequences, for consistency. Interestingly, we found substantial variation in the responses of both plants and butterflies, and no plant showed the same, elevated response to oviposition that we saw in *A. thaliana* (Fig. 4b, c). To investigate which, if any, components of the defense network were conserved among plants or among butterflies, we extracted and compared common differentially expressed genes across the four systems. Among plants, common gene families that were differentially expressed in leaves after oviposition were protein kinases, proteases, heat shock proteins, esterases, myb transcription factors and nac transcription factors (Supplementary data 5), all of which have previously been implicated in defense against pathogens^33,34^. During larval feeding, leaves of the four plant species shared protein kinases, proteases, heat shock proteins, esterases, myb transcription factors, nac transcription factors, and in addition, glutathione s-transerfases, cytochrome P450s, xyloglucan endotransglycosylases, dof zinc finger proteins and wrky transcription factors (Supplementary data 6). Based on protein alignments, we identified a small set of true orthologs among the differentially expressed genes, 11 in total, which were shared across all plant species. Leaves after oviposition did not have any shared orthologs but the larval feeding treatments had 4 or 5 orthologs each (Supplementary data 7), the majority of which have putative roles in defense.

**Figure 4:**
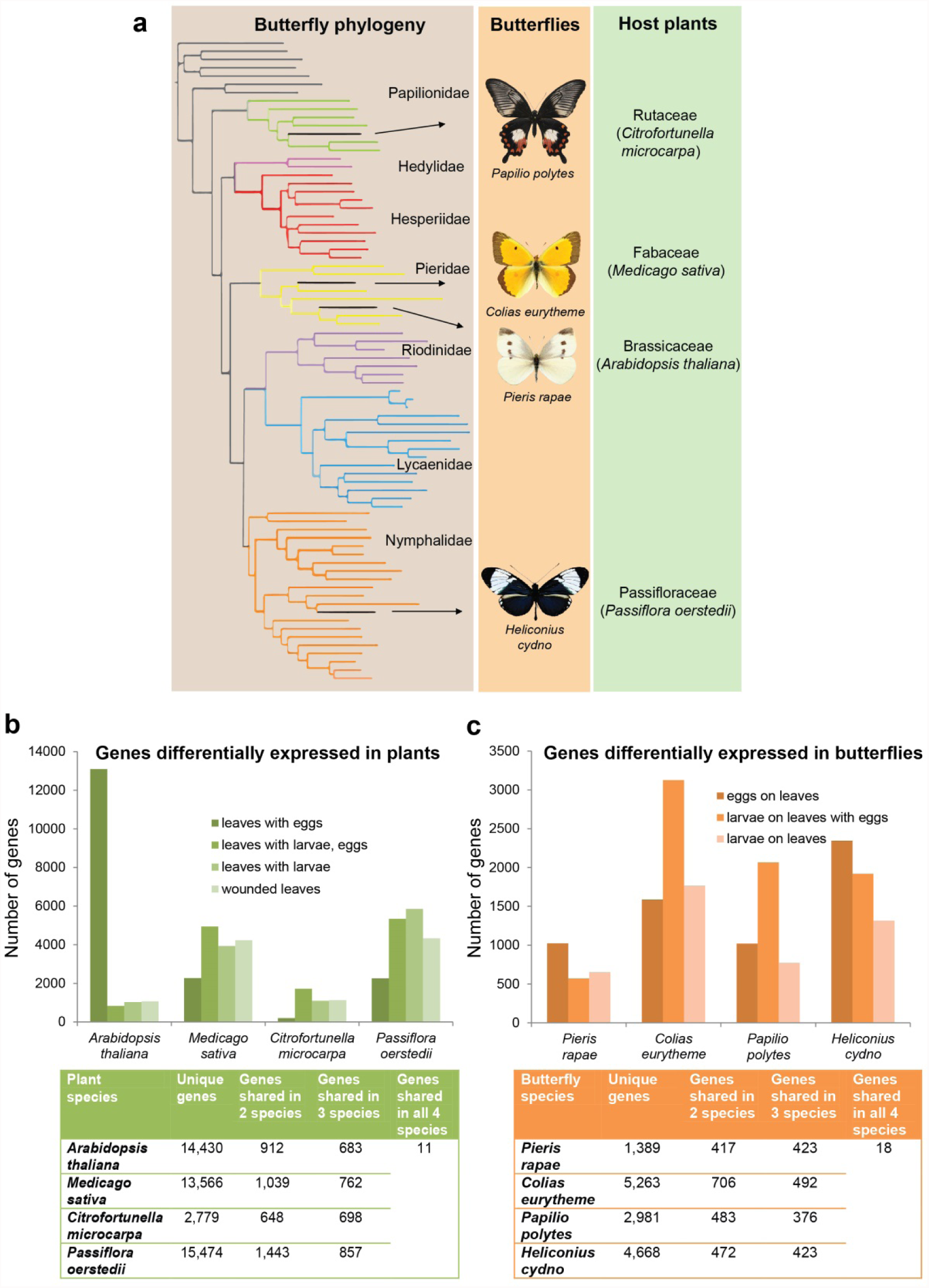
Comparative transcriptomics of herbivory across four diverse butterflies and their host-plants. (a) Butterfly phylogeny, adapted from ref. 79, displaying the four butterfly species included in the comparative transcriptomics analysis and their respective host-plants. (b) Number of differentially expressed genes across treatments for each host-plant species (above), and counts of differentially expressed genes that are unique or shared across species (below). (c) Number of differentially expressed genes across treatments for each butterfly species (above), and counts of differentially expressed genes that are unique or shared across species (below).

Among butterfly species, the common gene families that were differentially expressed in eggs 72 h following oviposition were proteases, heat shock proteins, esterases, chemosensory proteins and cuticular proteins (Supplementary data 8), all of which have previously been implicated in detoxification pathways^35,36^. In larvae feeding on plants, the genes differentially expressed belonged to gene families such as chitinases, proteases, cuticular proteins, lipases and Osiris (Supplementary data 9). There was a small set of differentially expressed genes, 17 in total, that were true orthologs shared among all butterfly species (Supplementary data 10). Among this set of conserved genes, *Osiris 9* (*Osi9*) stood out because it was up-regulated in all four butterfly species specifically in larvae feeding on leaves with previous exposure to eggs. *Osi9* is a transmembrane protein that is a member of the 24 gene *Osiris* family that is unique to insects 37. The function of *Osiris* genes has been mysterious^37,38^ but the *Osiris* gene cluster was recently associated with detoxification of the *Morinda citrifolia* host-plant in both *Drosophila sechellia*^39,40^ and *Drosophila yakuba*^41^. Previous work has shown that Lepidoptera appear to have multiple copies of *Osi9*^37^ and we detected multiple paralogs of *Osi9* in the *P. rapae* genome and across our assembled transcriptomes (Supplementary Fig. 3). However, the copy of *Osi9* that was up-regulated in larvae on all four butterfly species was the same, *Osiris 9E* (Supplementary Fig. 3). To further explore the potential role of *Osi9E* in butterflies, we surveyed spatial and temporal patterns of *Osi9E* expression in the four butterfly species and found elevated expression during larval stages, as opposed to pupae and adult, and expression specifically in the larval gut (Supplementary Figs. 4 and 5). These results suggest *Osi9E* expression is up-regulated in response to contact with host-plant tissue.

## Discussion

Butterflies and their larval host-plants provide a historically significant example of co-evolution and important prior research has explored the molecular genetic basis of this ubiquitous ecological interaction^5-8,15,16,30,31^. Here we used two genome-wide approaches, genome-wide association studies and transcriptomics, to characterize the genetic basis of herbivory in butterflies and plants simultaneously. Our genome-wide association studies uncovered a relatively small number of well-supported herbivory genes in both *A. thaliana* and *P. rapae*. The *A. thaliana* GWAS yielded 12 genes that contained three or more associated variants, eight of which we were able to validate with knock-out lines, and this gene set contained both established and novel plant defense genes. *P. rapae* GWAS revealed just one strongly associated region of the genome that contained only seven genes. One of these genes, *Glyoxalase 1*, stands out as a particularly good candidate gene because it utilizes glutathione to detoxify methylglyoxal, a toxic by-product of cellular metabolism. Glutathione is a defensive compound in plants: glutathione concentrations increase in plants during oxidative stress^42^^-^^45^ and it has been shown to be involved in defense against pathogens^46^ and insect feeding^47^. Furthermore, glutathione levels are known to vary among *Arabidopsis* accessions^48^ and we found natural sequence variation in the GST genes in our mapping panel (Supplementary data 11). It is intriguing to consider the possibility that *P. rapae* may somehow be using the glyoxalase pathway to detoxify host-plant glutathione. A second *P. rapae* GWA peak, while not statistically significant, was prominent and contained just a single gene, *peroxidasin*. *Peroxidasin* is critical for basement membrane integrity and tissue development, and this gene has been implicated in phagocytosis and defense^22,23^. The combined *P. rapae* GWA results suggest that we may have identified genes related to host-plant detoxification and/or larval metabolism and growth.

Previous research on *Pieris rapae* and related butterflies has identified the nitrile-specifier protein (NSP) as a critical component in host-plant detoxification^7^. The origin of NSP appears to be the key innovation that allowed the ancestor of the butterfly subfamily Pierinae to colonize and detoxify host-plants in the order Brassicales, all of which produce glucosinolates (i.e., mustard oils)^15,16^. Prior to our experiments, we hypothesized that genetic variation at NSP may also influence detoxification capacity in contemporary populations and that we may see a strong association with NSP in our butterfly GWA study. This was not the case. Furthermore, NSP was not differentially expressed in our transcriptomic experiments with *P. rapae*, although it was expressed in all larvae. In contrast, glucosinolate pathway genes did emerge in the *Arabidopsis* GWAS and many were differentially expressed in our transcriptomic studies, suggesting that glucosinolate defense remains an active front in this co-evolutionary arms race, at least on the plant side of the interaction.

Expanding our analysis to four diverse butterfly/plant systems, we found that over evolutionary time, the molecular dynamics of the plant-insect interaction change dramatically. The number, timing, and identity of genes expressed in plants and butterflies throughout the plant/insect interaction differed considerably across the four systems we studied. This finding highlights that the specific genetic underpinnings of herbivore-plant dynamics derived from a model system like *Arabidopsis* may not always be generalizable to other systems. However, such a result is also expected because the process of co-evolution should drive each system along a very different evolutionary trajectory, especially over the long times scales that separate the butterfly and plant species we are studying. What is surprising, then, is that we did identify a core set of orthologous genes that were differentially expressed in response to herbivory in all butterfly/plant systems. A total of 11 orthologs were differentially expressed in all plant species and 17 in butterflies, but this summary belies further complexity as a number of these genes, while all differentially expressed, were regulated in opposite directions among systems, being upregulated in one species and downregulated in another. One gene that stood out among the conserved orthologs in butterflies was *Osiris 9E*, both because it was consistently expressed at high levels in larvae of all four butterfly species and because *Osiris* genes were recently implicated in host-plant detoxification in *Drosophila*^39-41^. Furthermore, *Osiris 9* and a number of other *Osiris* genes have been shown to be differentially expressed in the larvae of the fly *Scaptomyza flava* in response to feeding on glucosinolates^49^. These results suggest that *Osiris* genes are ancient players in insect/plant interactions.

As a whole, our results provide a comprehensive portrait of the molecular genetic dynamics mediating insect/host interactions. Importantly, we are able to not only corroborate previous findings related to the identity of particular defense genes and pathways in *Arabidopsis*, but also expand this to other plants as well as the insect herbivores. The results from *Pieris* and other butterflies pinpoint specific genes that appear important for herbivory and these results further hint at even deeper evolutionary ties among herbivorous insects as a whole. Important next steps will be to functionally characterize these genes and identify the molecular and cellular mechanisms by which they impact herbivory. A complete understanding of the molecular genetic basis of the pervasive, antagonistic relationship between caterpillars and their host-plants promises to inform our understanding of ecology, evolution, and human agriculture.

## Methods

### Genome assembly and annotation

Genomic DNA was isolated from single, seventh-generation inbred female *Pieris rapae* pupae, using ethanol precipitation^50^. This inbred line of P. rapae was established using a singly-mated wild female caught in August 2013 near Rochester, Pennsylvania, USA (+40° 44’ 45”, -80° 9’ 45”). Three Illumina libraries were prepared, one PCR-free DNA library (180bp) and two mate-pair libraries (3kb and 7kb).The 180bp library was sequenced in two lanes and the two mate-pair libraries were sequenced in one lane each on Illumina HiSeq High Output mode, PE 100bp. Genomic DNA was isolated as described above from another inbred *P. rapae* pupae, a sibling of the first sample described above, and a 40kb mate pair library was constructed and sequenced by using a Lucigen NxSeq 40 kbp Mate-Pair Cloning Kit. For the final scaffolding step variable insert size libraries of 100bp – 100,000bp, using DNA from a third sibling, were generated using the Chicago and HiRise methods^51^ and these were sequenced by Centrillion Biosciences Inc. (Palo Alto, CA, USA), Illumina HiSeq High Output mode, PE100bp.

Genome size was estimated at 280 Mbp from unique k-mer distribution of the raw data using Jellyfish v2.1.3^52^ and a custom R script. For the 3kb and 7kb libraries, Nextclip (version 0.8)^53^ was used to look for the absence of linker sequence in either read in a pair and discard those reads as potential contamination of non-mate pair sequence. All read sets were then quality filtered, the ends trimmed of adapters and low quality bases, and screened of common contaminants using bbduk v34.94 (https://github.com/BioInfoTools/BBMap/blob/master/sh/bbduk.sh). For contig generation and scaffolding, the 180bp, 3kb, and 7kb reads were assembled using AllpathsLG (version 50960)^54^. The best assembly was obtained by using a random subset of 56 (33%) million reads from the initial 3kb and 7kb libraries with the full set of 162 million reads from the 180bp library, more input data resulted in reduced performance and quality of assembly. AllpathsLG was run with haploidify = true option to compensate for the high degree of heterozygosity present in the *P. rapae* data. The assembly was composed of 347 million bases contained in 8,113 scaffolds with a N50 of 141,106 bp. Complete conserved single copy ortholog content was assessed at 87% by CEGMA (version 2.5)^55^. A second scaffolding step using SSPACE v2^56^ and the 3kb, 7kb, and 40kb libraries together brought up the assembly size and the N50 to143, 392 bp. A final scaffolding step was undertaken by Dovetail genomics using the custom library and the HiRise scaffolding pipeline^51^, which improved the N50 to 3,706,409 bp. Chromosomal relationships of scaffolds were inferred from alignment using LAST (version 714)^57^ to the chromosomal structure of a closely related species *Pieris napi*^58^, and validated with scaffold junction spanning mate pair and syntenic blocks (Supplementary Fig. 1). The final *P. rapae* assembly contained 323,179,347 bp in 25 chromosomes and 2,747 unplaced scaffolds and a N50 of 11,535,178 bp.

The genome annotation involved the following pipeline: a) Collection of reference proteins from Uniprot database^59^ and assembly of high-confidence transcript sequences from previously published RNA-seq data^60^ using tophat2 (version 2.0.9)^61,62^ and the cufflinks package (version 2.2.1)^63^, b) Modelling of repeat sequences to mask the genome using RepeatMasker package (4.0.3) (Smit, AFA, Hubley, R & Green, P. RepeatMasker Open-3.0.1996-2010 <http://www.repeatmasker.org>) and RepeatModeler package (1.0. 8) (Smit, AFA, Hubley, R. RepeatModeler Open-1.0.2008-2015 <http://www.repeatmasker.org>), c) Evidence-based gene build to generate training models for ab-initio gene finders using the Maker package (version 2.31-6)^64^, d) Manual curation of gene models and training of the Augustus gene finder (version 2.7)^65^, e) Re-annotation of the evidence-based annotation using ab-initio predictions, and finally, f) Functional annotation of the refined gene build using Blast matches against Uniprot/Swissprot and results from InterproScan, condensed and reconciled using ANNotation Information Extractor(Annie)( Tate, R., Hall, B., DeRego, T., & Geib, S. (2014). Annie: the ANNotation Information Extractor (Version 1.0)).

### Sample collection and data analysis for GWAS

Host-plant GWAS: The growth chamber conditions for growing *Pieris rapae* and *Arabidopsis thaliana* were 23°C day/21°C night and 60% relative humidity on a 16 hour photoperiod. Three replicates of each of the 96 accessions of *Arabidopsis thaliana* that are listed in Supplementary Table 1 were grown until they were almost ready to bolt. After taking a picture of each plant, two 5 day old, lab-grown *P. rapae* larvae were weighed and placed on each plant and then the plant was enclosed in a plastic sleeve bag. After 72 h, the larvae were weighed and a new picture of the plant was taken in order to record plant surface area eaten by the larvae. The weight gained by the larvae and the total surface area eaten were calculated and used as the phenotype data for GWAS. The SNP information for the 96 accessions from the 250K SNP data^66^ and the phenotype data were fit into a multivariate linear mixed model in GEMMA (version 0.94.1) for association studies^67^. The pipeline involves converting the SNP file into a PLINK binary PED file and generating a relatedness matrix file using default parameters. The average initial weight of the larvae was used as a covariate for the analysis. For the functional validation of the candidate genes, knock out T-DNA mutants from SALK and wild type plants were grown and assayed as described above.

Herbivore GWAS: 96 *P. rapae* females were collected from various locations across the US Midwest during June-July 2014 and raised in the lab green house. These butterflies included 57 from around the University of Chicago campus, 19 from Schaumburg, IL, and 16 from North Dakota, 3 from downtown Chicago, and 1 from Carolina Biological Supply. The growth chamber conditions for growing *P. rapae* and *A. thaliana* were 23°C day/21°C night and 65% relative humidity on a 16 hour photoperiod. Eggs were collected from each female and two 5 day old larvae from each family were weighed and placed together on a Col-0 *A. thaliana* plant and this was performed in triplicate. Leaf area eaten and weight gain phenotypes were assayed as described for the *Arabidopsis* GWAS. For genotyping, we selected one random GWAS larva from each family (a total of 96 larvae) and sequenced the genome of each sample at 5X coverage and analyzed using the *P. rapae* genome assembly. Genomic DNA was extracted from skin tissue of the larvae using the VDRC Drosophila genomic extraction (https://stockcenter.vdrc.at/images/downloads/GoodQualityGenomicDNA.pdf) and sequencing libraries were prepared using KAPA Hyper Prep Kits (KR0961 – v1.14). Barcoded libraries were pooled and sequenced on an Illumina HiSeq 2500 to generate paired-end 100 bp data. After sequencing, PE100 bp Illumina reads were trimmed using Trimmotatic (version 0.36)^68^, aligned to the *P. rapae* reference genome using bowtie2, using a –very-sensitive-local option, and the aligned SAM files were prepared for calling SNPs using the default parameters of PICARD tools (version 1. 141)(http://broadinstitute.github.io/picard). Finally, UnifiedGenotyper, a variant calling program in GATK (version 3.4)^69,70^ was used to call SNPs. The number of reads generated and the number of SNPs called per sample is listed in Supplementary Table 2.The SNP information for the 96 larvae (a total of 18,603,675 SNPs) and the phenotype data were fit into a multivariate linear mixed model in GEMMA (version 0.94.1) for association studies^67^. The pipeline involves converting the SNP file into a PLINK binary PED file and generating a relatedness matrix file using default parameters. Again, the average initial weight of the larvae was used as a covariate for the analysis. The statistical significance for both the *Arabidopsis* and *Pieris* GWAS was determined by simpleM, a multiple testing correction method for genetic association studies using correlated SNPs^71^.

### *Arabidopsis* and *Pieris* RNA-seq

*Pieris rapae* was raised on *Arabidopsis thaliana* at 23°C and 65% relative humidity on a 16 hour photoperiod. Samples at different time points were collected from both the plant and insect in TRIzol Reagent. The time points are listed in Supplemental Table 3. RNA was extracted using TRIzol Reagent and the RNA-seq libraries were generated using an Illumina TruSeq Stranded Total RNA Kit with Ribo-Zero Plant for *Arabidopsis* and Illumina TruSeq Stranded mRNA Library Prep Kit for *Pieris.* These libraries were sequenced on an Illumina HiSeq 2500 to generate SR50bp reads. All the SR50bp reads were processed with the Trimmomatic (version 0.36)^68^ QC pipeline. The trimmed SR50bp reads were aligned to their respective genomes using STAR (version 2.4.2)^72^ software. The Cuffdiff pipeline (version 2.2.1)^63^ was used to call the differentially expressed genes at using an FDR-corrected p-value of 0.001. The differentially expressed set of genes in *Arabidopsis thaliana* and *Pieris rapae* were annotated using the annotations available for their respective genomes.

### Comparative transcriptome analysis

Each butterfly species was reared on its natural host-plant under favorable environmental conditions in the laboratory greenhouse. *Pieris rapae* was raised on *Arabidopsis thaliana* at 23°C and 65% relative humidity on a 16 hour photoperiod. *Colias eurytheme* was raised on *Medicago sativa* at 26°C and 65% relative humidity on a 16 hour photoperiod. *Papilio polytes* was raised on *Citrofortunella microcarpa* at 26°C and 65% humidity on a 16 hour photoperiod. *Heliconius cydno* was raised on *Passiflora oerstedii* at 26°C and 65% relative humidity for a 13 hour photoperiod. Samples at different time points were collected from both plants and herbivores in TRIzol Reagent. These are the same time points as those listed in Supplemental Table 3. RNA was extracted using TRIzol Reagent, and RNA-seq libraries were generated using Illumina TruSeq Stranded Total RNA Kit with Ribo-Zero Plant for the four plant species and Illumina TruSeq Stranded mRNA Library Prep Kit for the four butterfly species. These libraries were sequenced on an Illumina HiSeq 2500 to generate SR50bp reads. In addition, one library from each of the host-plant species and two libraries from of each butterfly species (the egg and a larval stage) were sequenced with PE100bp in order to generate a reference transcriptome for each species. The number of reads generated from each library is listed in Supplemental Table 4.

The SR50bp and PE100bp reads were processed with the Trimmomatic QC pipeline (version 0.36)^68^. All trimmed PE100bp reads and the trimmed SR50bp reads were assembled together for each host-plant and herbivore system to generate a reference transcriptome. After excluding different isoforms, the number of unique coding sequences found in each plant species was: *A. thaliana*, 18,105; *M. sativa*, 43,843; *C. microcarpa*, 20,367; and *P. oerstedii*, 22,073. The number of unique coding sequences in each butterfly species was: *P. rapae*, 12,037; *C. eurytheme*, 16,277; *P. polytes*, 15,176; and *H. cydno*, 19,914. Trimmed SR50bp reads from each species were aligned to their respective reference transcriptome and differential expression analyses were performed with the Trinity package (version r20140717)^73^, which includes bowtie2 for alignment, RSEM for transcript abundance and edgeR for different expression analysis. Default parameters and an FDR-adjusted p-value of 0.001 were used for analyses. The differentially expressed set of genes from all four host-herbivore systems were annotated using Blast2GO (version 3.3)^74^ using default settings. The annotated genes were manually parsed to find the common gene families. To detect true orthologs that were differentially expressed across all plants or all butterflies, we analyzed the protein sequences of the differentially expressed genes with Proteinortho (version 5.11)^75,76^ using a p-value of 0.001.

### *Osiris 9* paralog detection

All *Osiris 9* protein sequences from *Bombyx mori*, *Drosophila melanogaster* and *Danaus plexippus* were extracted from the *Osiris* protein database^37^. Using them as the query, the homologous sequences were extracted from the *de novo* transcriptomes of all the four butterfly species and the *P. rapae* genome using tblastn similarity searches. A multiple sequence alignment of all extracted *Osiris 9* protein sequences, including the *Osiris 11* and *Osiris 14* from *B. mori*, was generated using MAFFT v6.847b^77^ with the L-INS-i algorithm. A phylogenetic tree was reconstructed using FastTree (version 2.1.10), which infers approximately maximum likelihood phylogenies from alignments of protein sequences^78^. All the default options and the JTT+CAT model were used for tree inference.

### *Osiris 9E* gene spatial and temporal assay

For the temporal assay of *Osiris 9E* expression in butterflies, whole body tissue samples were collected at ten stages of development for all four butterfly species, except for adults where wings and eyes were excluded. Multiple individuals were pooled at early instar stages (24 hours to 72 hour instar) because of their small size. For the rest of the stages, a single individual was used for each RNA extraction. The developmental stages were: 24 hour larvae, 48 hour larvae, 72 hour larvae, 2nd instar larva, 3rd instar larva, 4th instar larva, 5th instar larva, pre-pupa, 72 hour pupa, and adult. For the spatial assay of *Osiris 9E* expression in larvae, the head, foregut, midgut, and skin were dissected out for each individual. RNA was extracted using TRIzol Reagent, and cDNA for each sample was synthesized using a High-Capacity cDNA Reverse Transcription Kit (Applied Biosystems) using 500 ng/ul of RNA. RT-qPCR for the temporal and spatial samples of each species was performed using SsoAdvanced SYBR Green Supermix (Bio-Rad) and a CFX96 Optical Reaction Module thermocycler (Bio-Rad). *Elongation Factor 1 alpha* ( *EF1α*) served as a normalizing gene to measure the relative expression of *Osiris 9E* in each sample for all four butterfly systems, the primer sequences for each species are in Supplemental Table 5.

## Acknowledgments

We thank E. Westerman, R. Marquez, L. Southcott, A. Russell and G. Garcia for assistance with experiments and D. Samac for generously providing *Medicago sativa* seeds. N. Saleh helped with generating the inbred *P. rapae* material for the sequencing the initial genome assembly. We also thank Cristina Sahagun for assisting with the butterfly photography. This project was funded by NIH grant GM108626 and NSF grant IOS-1452648 to MRK, as well as the Pew Charitable Trust and Neubauer funds from the University of Chicago and University of Pittsburgh.

## Author Contributions

SN designed the project, collected and analyzed data for the GWAS, RNA-seq and comparative transcriptome studies, and co-wrote the manuscript. JH and CW sequenced and assembled the *Pieris rapae* genome. KD, CS and WZ helped collect and raise butterflies and contributed to the GWAS experiments. KD analyzed temporal and spatial expression patterns for *Osi9E* expression. NM generated the inbred *P. rapae* line used for genome sequencing, and NM, CM and NC generated the RNA-seq data used for *P. rapae* genome annotation. ES-R collected the North Dakota *P. rapae* individuals for GWAS. JB provided *Arabidopsis* accessions for GWAS. MRK designed and directed the project, and wrote the manuscript. All authors contributed to editing the manuscript.

## Materials & Correspondence

Correspondence to: M. R. Kronforst

## Figure legends

**Supplementary Figure 1:**
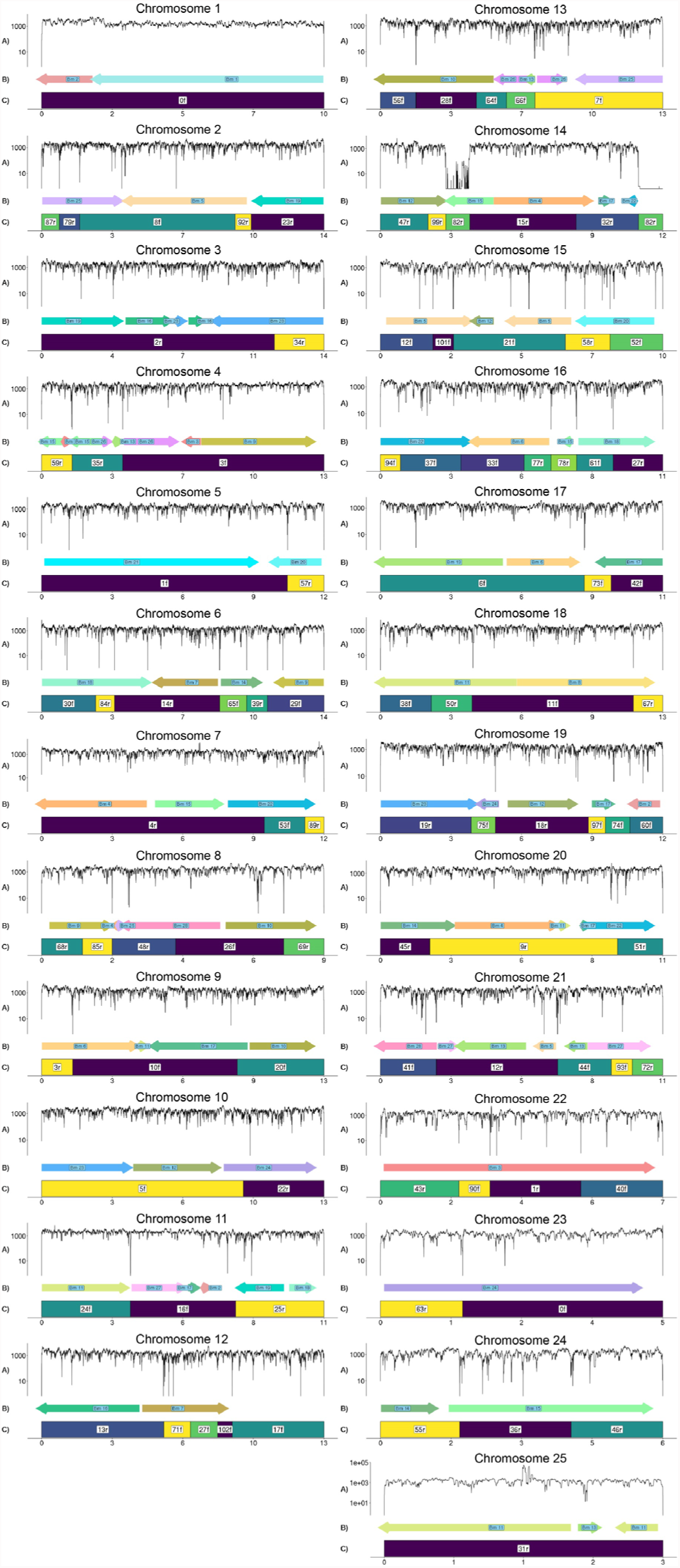
Chromosomal assembly of butterfly *Pieris rapae* genome. (a) Read pairs from the 3 kbp, 7 kbp, and 40 kbp insert size mate pair libraries were mapped to the *P. rapae* chromosomes and the number of times a position was flanked by read pair mapped in the proper orientation with a mapq > 20 for both reads was counted. A count of zero occurs at a position where no mate pairs span which can be caused by either a misassembly or a high density repetitive sequence. (b) Orthologous genes from *Bombyx mori* occur in blocks along the *P. rapae* chromosomes and their *B. mori* chromosome of origin and orientation are shown above (c) the number and direction, forward or reverse, of the component scaffold used to construct the chromosome. A chromosome misassembly is less likely to occur within a scaffold, within a syntenic block, or where mate pair spanning is high. Taken together we regard a scaffold join that is spanned by an average number of mate pairs and a syntenic block from *B. mori* to be correct.

**Supplementary Figure 2:**
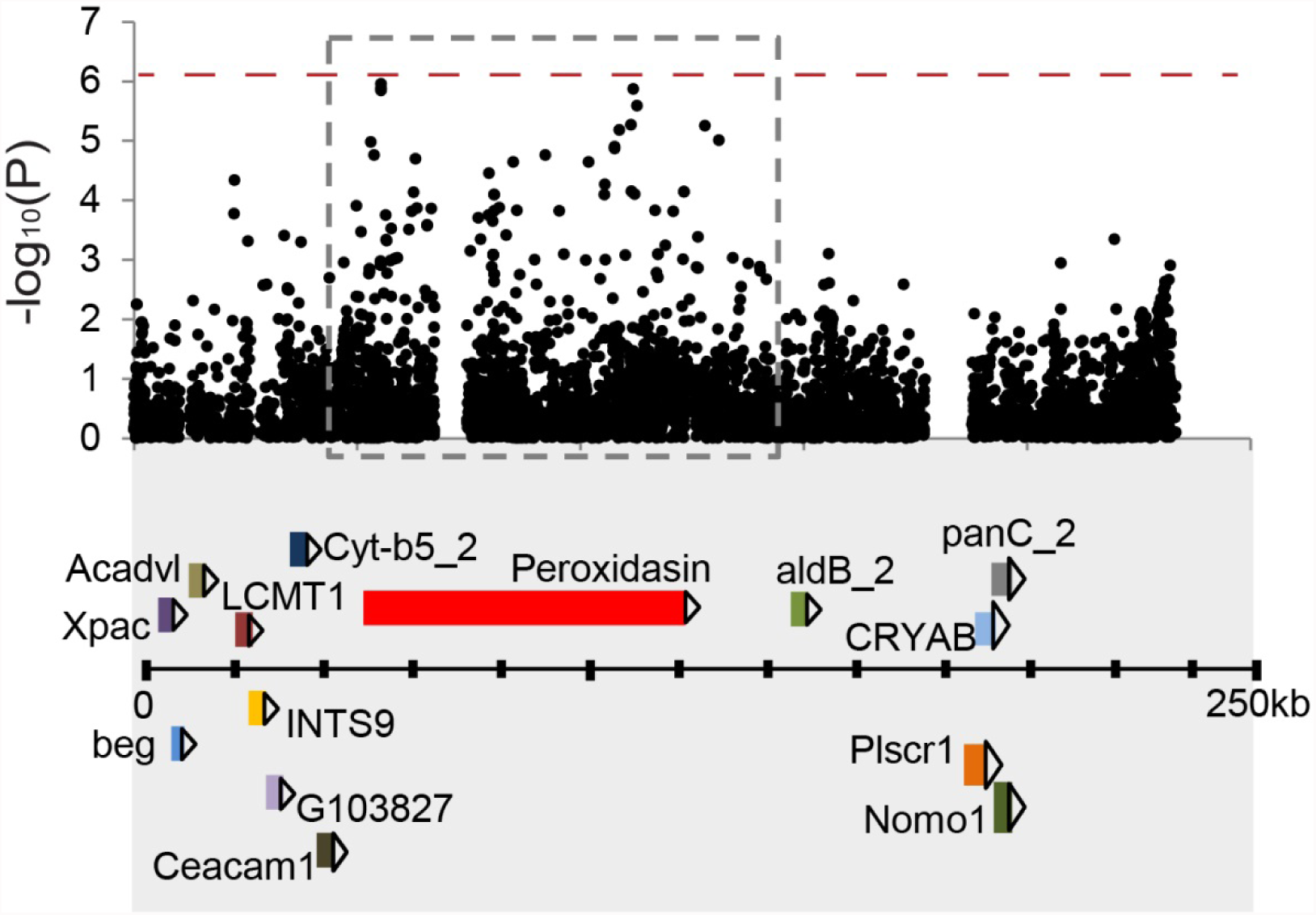
Zoom-in on the chromosome 15 region (boxed) associated with herbivory in the *P. rapae* genome. A gene map below shows the underlying genes found in the associated and the adjoining regions.

**Supplementary Figure 3:**
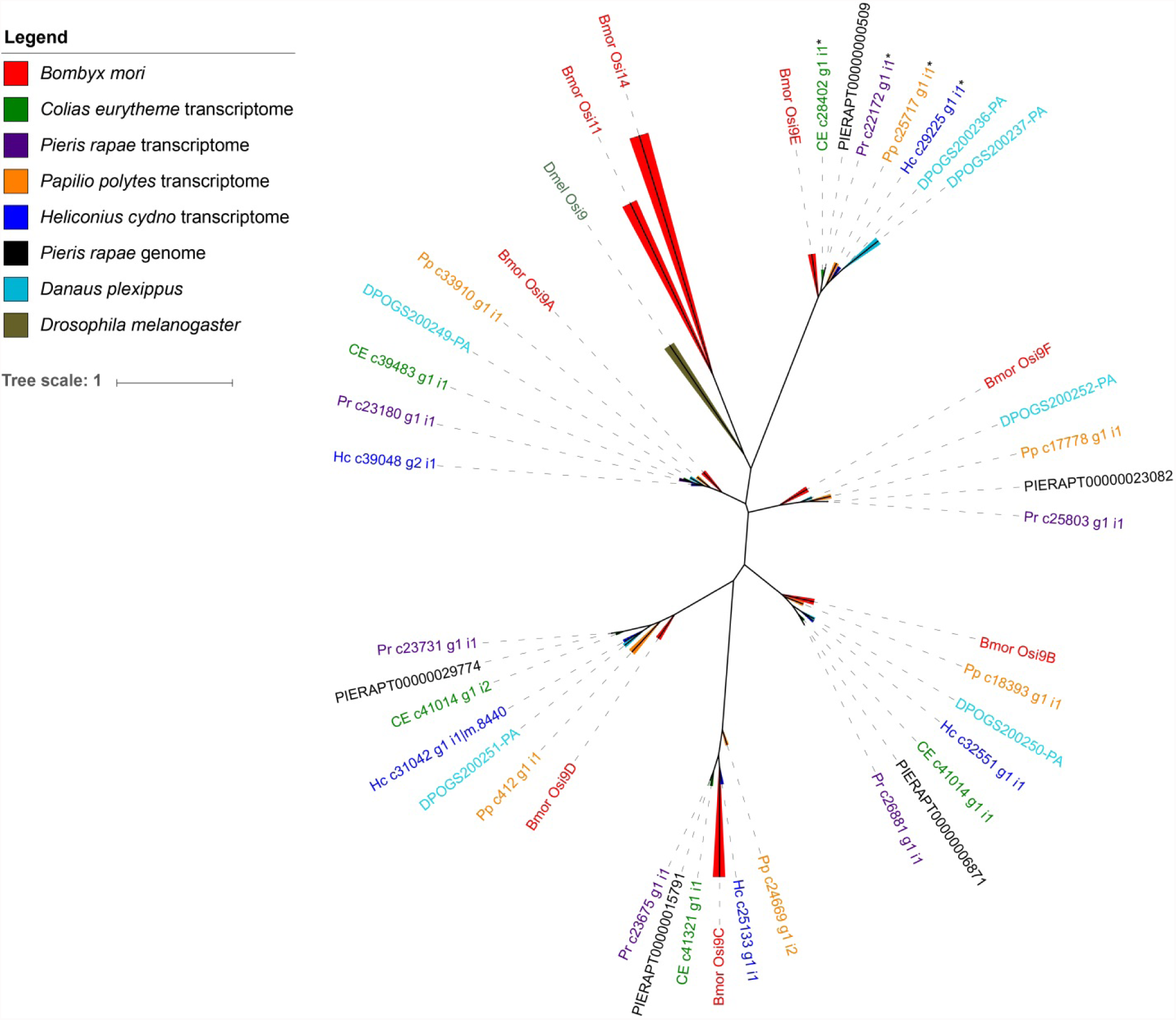
Tree of *Osiris 9* protein sequences extracted from the *Pieris rapae* genome and each of the four assembled transcriptomes—*Colias eurytheme*, *Pieris rapae*, *Papilio polytes*, and *Heliconius cydno*. These are combined with *Osiris 9* protein sequences from *Drosophila melanogaster*, *Bombyx mori*, and *Danaus plexippus*, as well as *Osiris 11* and *Osiris 14* from *B. mori*. While *Osi9* is clearly a multi-copy gene across the Lepidoptera, we found one copy to be up-regulated in larvae of all four butterfly species when feeding on leaves with prior exposure to eggs (Supplementary data 10). These genes are marked with an asterisk and the *Osi9* tree shows that they are all *Osi9E*.

**Supplementary Figure 4:**
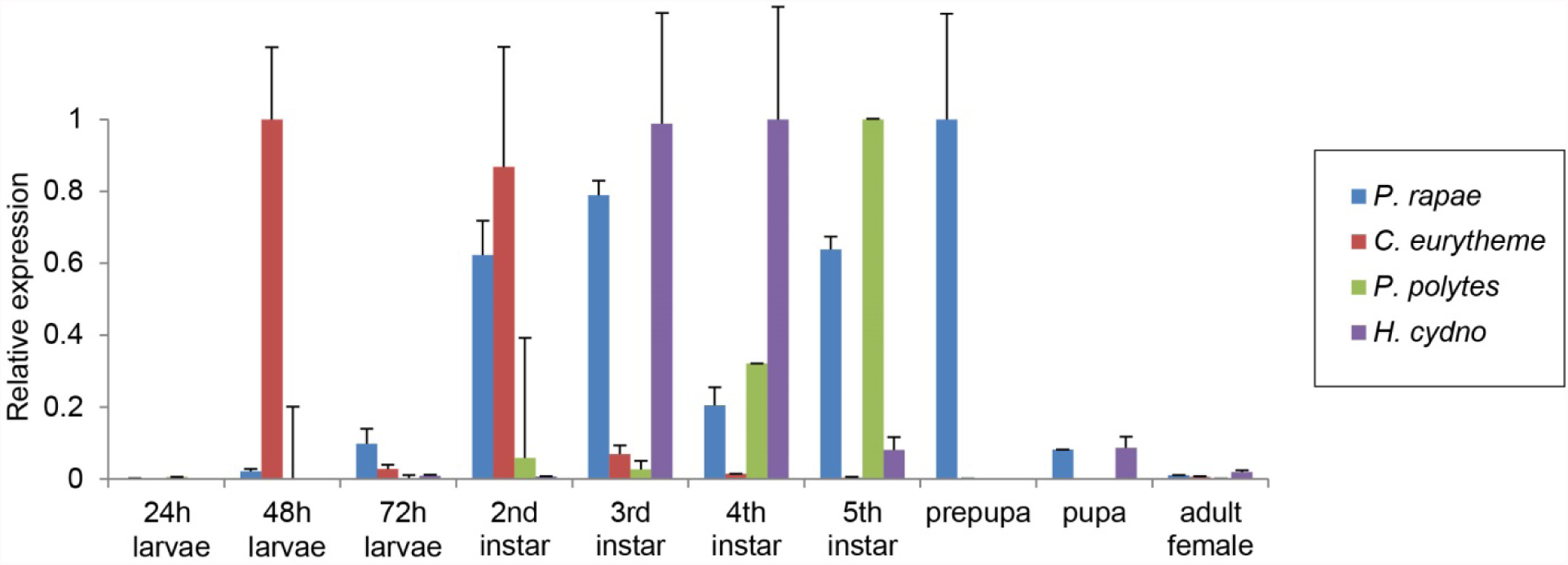
Temporal expression patterns of *Osiris 9E* with positive standard error of the mean for all the three biological replicates in a given developmental stage. Developmental stages are represented along the x-axis with relative average fold change levels of *Osi9E* along the y-axis.

**Supplementary Figure 5:**
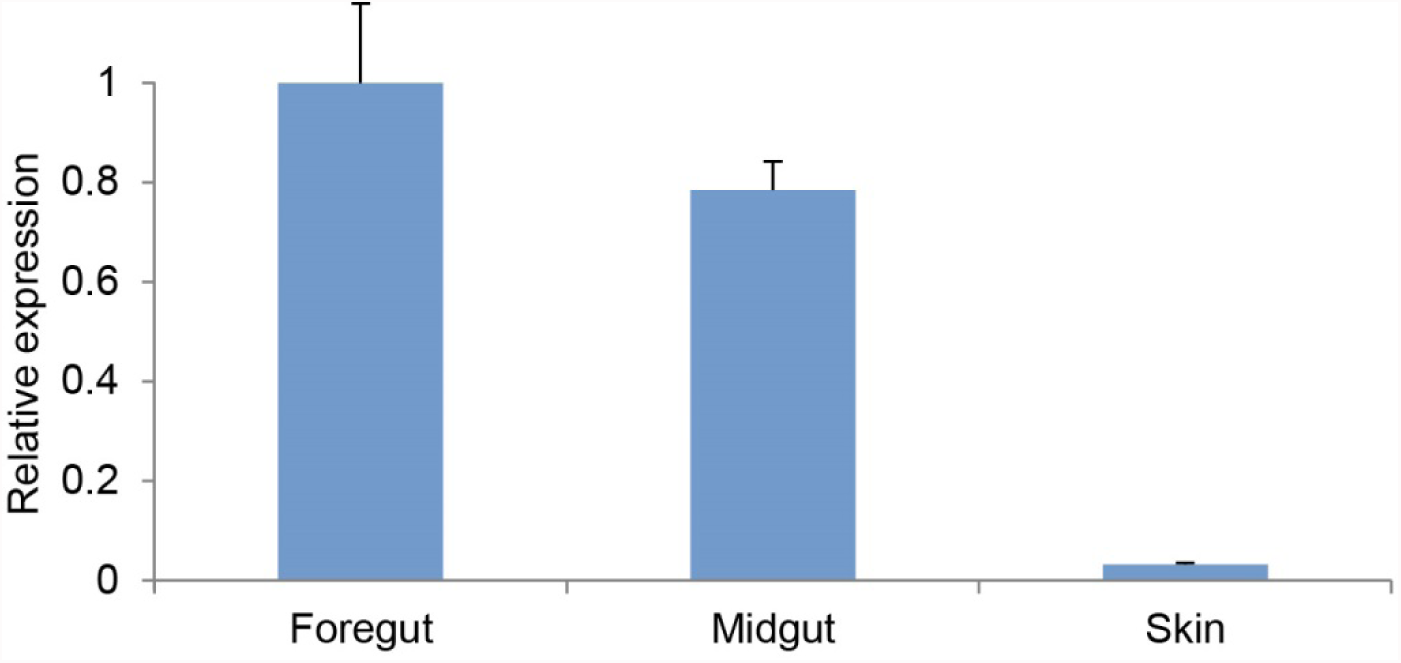
Spatial expression patterns of *Osiris 9E* with positive standard error of the mean for all the three biological replicates in the larval 3rd instar of *P. rapae*. Various parts of the body are represented along the x-axis with relative average fold change levels of *Osi9E* along the y-axis.

**Supplementary Table 1.** List of 96 *A.thaliana* accessions used in Arabidopsis GWAS.

| **Accession name** | **country** | **Stockparent** | **collector** | **Collection date** | **country** | **Latitude** | **Longitude** | **Site** |
| --- | --- | --- | --- | --- | --- | --- | --- | --- |
| Ag-0 | France | CS22630 | Albert Kranz | NA | France | 45 | 1.3 | Ag |
| An-1 | Belgium | CS22626 | Albert Kranz | NA | Belgium | 51.2167 | 4.4 | An |
| Bay-0 | Germany | CS22633 | Albert Kranz | NA | Germany | 49 | 11 | Bay |
| Bil-5 | Sweden | CS22578 | Magnus Nordborg | 6/25/2000 | Sweden | 63.324 | 18.484 | Bil |
| Bil-7 | Sweden | CS22579 | Magnus Nordborg | 6/25/2000 | Sweden | 63.324 | 18.484 | Bil |
| Bor-1 | Czech | CS22590 | Jirina Relichová | 5/31/2000 | Czech Republic | 49.4013 | 16.2326 | Bor |
| Bor-4 | Czech | CS22591 | Jirina Relichová | 5/31/2000 | Czech Republic | 49.4013 | 16.2326 | Bor |
| Br-0 | Czech | CS22628 | Albert Kranz | NA | Czech Republic | 49.2 | 16.6166 | Br |
| Bur-0 | Ireland | CS22656 | Albert Kranz | NA | Ireland | 54.1 | -6.2 | Bur |
| C24 | Portugal | CS22620 | Brigitte Damm | NA | Portugal | 40.2077 | -8.42639 | Co |
| CIBC-17 | UK | CS22603 | Mick Crawley | 5/21/1993 | United Kingdom | 51.4083 | -0.6383 | CIBC |
| CIBC-5 | UK | CS22602 | Mick Crawley | 5/21/1993 | United Kingdom | 51.4083 | -0.6383 | CIBC |
| Col-0 | USA | CS22625 | Albert Kranz | NA | United States of America | 38.3 | -92.3 | Col |
| Ct-1 | Italy | CS22639 | Albert Kranz | NA | Italy | 37.3 | 15 | Ct |
| Cvi-0 | Cape Verde | CS22614 | Albert Kranz | NA | Cape Verde | 15.1111 | -23.6167 | Cvi |
| Eden-1 | Sweden | CS22572 | Magnus Nordborg | 6/25/2000 | Sweden | 62.877 | 18.177 | Eden |
| Eden-2 | Sweden | CS22573 | Magnus Nordborg | 6/25/2000 | Sweden | 62.877 | 18.177 | Eden |
| Edi-0 | UK | CS22657 | Albert Kranz | NA | United Kingdom | 55.949444 | -3.160278 | Edi |
| Ei-2 | Germany | CS22616 | Albert Kranz | NA | Germany | 50.3 | 6.3 | Ei |
| Est-1 | Russia | CS22629 | Albert Kranz | NA | Russia | 58.3 | 25.3 | Est |
| Fäb-2 | Sweden | CS22576 | Magnus Nordborg | 6/25/2000 | Sweden | 63.0165 | 18.3174 | Fäb |
| Fäb-4 | Sweden | CS22577 | Magnus Nordborg | 6/25/2000 | Sweden | 63.0165 | 18.3174 | Fäb |
| Fei-0 | Portugal | CS22645 | Carlos Alonso-Blanco | NA | Portugal | 40.5 | -8.32 | Fei |
| Ga-0 | Germany | CS22634 | Albert Kranz | NA | Germany | 50.3 | 8 | Ga |
| Got-22 | Germany | CS22609 | Gerhard Röbbelen | NA | Germany | 51.5338 | 9.9355 | GOT |
| Got-7 | Germany | CS22608 | Gerhard Röbbelen | NA | Germany | 51.5338 | 9.9355 | GOT |
| Gu-0 | Germany | CS22617 | Albert Kranz | NA | Germany | 50.3 | 8 | Gu |
| Gy-0 | France | CS22631 | Albert Kranz | NA | France | 49 | 2 | Gy |
| HR-10 | UK | CS22597 | Mick Crawley | 5/21/1993 | United Kingdom | 51.4083 | -0.6383 | HR |
| HR-5 | UK | CS22596 | Mick<br>Crawley | 5/21/1993 | United<br>Kingdom | 51.4083 | -0.6383 | HR |
| Kas-2 | India | CS6751 | Shauna<br>Somerville | 1/1/1976 | India | 35 | 77 | Kas |
| Kin-0 | USA | CS22654 | Albert<br>Kranz | NA | United<br>States of<br>America | 44.46 | -85.37 | Kin |
| Kno-10 | USA | CS22566 | Joy<br>Bergelson | 34111 | United<br>States of<br>America | 41.2816 | -86.621 | Knox |
| Kno-18 | USA | CS22567 | Joy<br>Bergelson | 34111 | United<br>States of<br>America | 41.2816 | -86.621 | Knox |
| Kondara | Tajikistan | CS22651 | Igor Vizir | NA | Tajikistan | 38.48 | 68.49 | Kondara |
| Kz-1 | Kazakhstan | CS22606 | Ihsan Al-Shehbaz | 6/7/1994 | Kazakhstan | 49.5 | 73.1 | KZ |
| Kz-9 | Kazakhstan | CS22607 | Ihsan Al-Shehbaz | 6/7/1994 | Kazakhstan | 49.5 | 73.1 | KZ |
| Ler-1 | Germany | CS22618 | Eric Holub | NA | Germany | 47.984 | 10.8719 | Ler |
| LL-0 | Spain | CS22650 | Albert<br>Kranz | NA | Spain | 41.59 | 2.49 | Ll |
| Löv-1 | Sweden | CS22574 | Magnus Nordborg | 6/25/2000 | Sweden | 62.801 | 18.079 | Löv |
| Löv-5 | Sweden | CS22575 | Magnus Nordborg | 6/25/2000 | Sweden | 62.801 | 18.079 | Löv |
| Lp2-2 | Czech | CS22594 | Ivo Cetl | NA | Czech Republic | 49.38 | 16.81 | Lp2 |
| Lp2-6 | Czech | CS22595 | Ivo Cetl | NA | Czech Republic | 49.38 | 16.81 | Lp2 |
| Lz-0 | France | CS22615 | Albert Kranz | NA | France | 46 | 3.3 | Lz |
| Mr-0 | Italy | CS22640 | Albert Kranz | NA | Italy | 44.15 | 9.65 | Mr |
| Mrk-0 | Germany | CS22635 | Albert Kranz | NA | Germany | 49 | 9.3 | Mrk |
| Ms-0 | Russia | CS22655 | Albert Kranz | NA | Russia | 55.7522 | 37.6322 | Ms |
| Mt-0 | Libya | CS22642 | Albert Kranz | NA | Libya | 32.34 | 22.46 | Mt |
| Mz-0 | Germany | CS22636 | Albert Kranz | NA | Germany | 50.3 | 8.3 | Mz |
| N13 | Russia | CS22491 | Andy Savushkin | NA | Russia | 61.36 | 34.15 | Konchezero |
| Nd-1 | Switzerland | CS22619 | Eric Holub | NA | Switzerland | 50 | 10 | Nd |
| NFA-10 | UK | CS22599 | Mick<br>Crawley | 5/21/1993 | United<br>Kingdom | 51.4083 | -0.6383 | NFA |
| NFA-8 | UK | CS22598 | Mick<br>Crawley | 5/21/1993 | United<br>Kingdom | 51.4083 | -0.6383 | NFA |
| Nok-3 | Netherlands | CS22643 | Albert<br>Kranz | NA | Netherlands | 52.24 | 4.45 | Nok |
| ÖMö2-1 | Sweden | CS22584 | Magnus<br>Nordborg | 5/14/2000 | Sweden | 56.1481 | 15.821214 | ÖMö2 |
| ÖMö2-3 | Sweden | CS22585 | Magnus<br>Nordborg | 5/14/2000 | Sweden | 56.147363 | 15.814587 | ÖMö2 |
| Oy-0 | Norwegia | CS22658 | Albert<br>Kranz | NA | Norway | 60.385543 | 6.193019 | Oy |
| Pna-10 | USA | CS22571 | Joy<br>Bergelson | NA | United<br>States of<br>America | 42.0945 | -86.3253 | PNA |
| Pna-17 | USA | CS22570 | Joy<br>Bergelson | NA | United<br>States of<br>America | 42.0945 | -86.3253 | PNA |
| Pro-0 | Spain | CS22649 | Joy<br>Bergelson | NA | Spain | 43.25 | -6 | Pro |
| Pu2-23 | Czech | CS22593 | Ivo Cetl | NA | Czech Republic | 49.42 | 16.36 | Pu2 |
| Pu2-7 | Czech | CS22592 | Ivo Cetl | NA | Czech Republic | 49.42 | 16.36 | Pu2 |
| Ra-0 | France | CS22632 | Albert Kranz | NA | France | 46 | 3.3 | Ra |
| Ren-1 | France | CS22610 | Gerhard Röbbelen | NA | France | 48.5 | -1.41 | REN |
| Ren-11 | France | CS22611 | Gerhard Röbbelen | NA | France | 48.5 | -1.41 | REN |
| Rmx-A02 | USA | CS22568 | Joy Bergelson | NA | United States of America | 42.036 | -86.511 | RMX |
| Rmx-A180 | USA | CS22569 | Joy Bergelson | NA | United States of America | 42.036 | -86.511 | RMX |
| RRS-10 | USA | CS22565 | Joy Bergelson | NA | United States of America | 41.5609 | -86.4251 | RRS |
| RRS-7 | USA | CS22564 | Joy Bergelson | NA | United States of America | 41.5609 | -86.4251 | RRS |
| Se-0 | Spain | CS22646 | Albert Kranz | NA | Spain | 38.3333 | -3.53333 | Se |
| Shahdara | Tajikistan | CS22652 | Igor Vizir | NA | Tajikistan | 38.35 | 68.48 | Sha |
| Sorbo | Tajikistan | CS22653 | Igor Vizir | NA | Tajikistan | 38.35 | 68.48 | Sorbo |
| Spr1-2 | Sweden | CS22582 | Magnus Nordborg | 5/15/2000 | Sweden | 58.4168 | 14.1612 | Spr1 |
| Spr1-6 | Sweden | CS22583 | Magnus Nordborg | 5/15/2000 | Sweden | 58.4168 | 14.1612 | Spr1 |
| Sq-1 | UK | CS22600 | Mick Crawley | 5/21/1993 | United Kingdom | 51.4083 | -0.6383 | SQ |
| Sq-8 | UK | CS22601 | Mick Crawley | 5/21/1993 | United Kingdom | 51.4083 | -0.6383 | SQ |
| Tamm-2 | Finland | CS22604 | Outi Savolainen | NA | Finland | 60 | 23.5 | Tamm |
| Tamm-27 | Finland | CS22605 | Outi Savolainen | NA | Finland | 60 | 23.5 | Tamm |
| Ts-1 | Spain | CS22647 | Albert Kranz | NA | Spain | 41.7194 | 2.93056 | Ts |
| Ts-5 | Spain | CS22648 | Albert Kranz | NA | Spain | 41.7194 | 2.93056 | Ts |
| Tsu-1 | Japan | CS22641 | Eric Holub | NA | Japan | 34.43 | 136.31 | Tsu |
| Ull2-3 | Sweden | CS22587 | Magnus Nordborg | 5/14/2000 | Sweden | 56.0648 | 13.9707 | Ull2 |
| Ull2-5 | Sweden | CS22586 | Magnus Nordborg | 5/14/2000 | Sweden | 56.0648 | 13.9707 | Ull2 |
| Uod-1 | Austria | CS22612 | Marcus Koch | 6/23/2000 | Austria | 48.3 | 14.45 | Uod |
| Uod-7 | Austria | CS22613 | Marcus Koch | 6/23/2000 | Austria | 48.3 | 14.45 | Uod |
| Van-0 | Canada | CS22627 | Albert Kranz | NA | Canada | 49.2655 | -123.206 | Van |
| Vår2-1 | Sweden | CS22580 | Magnus Nordborg | 6/1/2000 | Sweden | 55.58 | 14.334 | V?r2 |
| Vår2-6 | Sweden | CS22581 | Magnus Nordborg | 6/1/2000 | Sweden | 55.58 | 14.334 | V?r2 |
| Wa-1 | Poland | CS22644 | Albert Kranz | NA | Poland | 52.3 | 21 | Wa |
| Wei-0 | Switzerland | CS22622 | Alan Slusarenko | NA | Switzerland | 47.25 | 8.26 | Wei |
| Ws-0 | Russia | CS22623 | Albert Kranz | NA | Russia | 52.3 | 30 | Ws |
| Ws-2 | Russia | CS22659 | Kenneth Feldmann | NA | Russia | 52.3 | 30 | Ws |
| Wt-5 | Germany | CS22637 | Albert<br>Kranz | NA | Germany | 52.3 | 9.3 | Wt |
| Yo-0 | USA | CS22624 | Albert<br>Kranz | NA | United<br>States of<br>America | 37.45 | -119.35 | Yo |
| Zdr-1 | Czech | CS22588 | Jirina<br>Relichová | 5/31/2000 | Czech<br>Republic | 49.3853 | 16.2544 | Zdr |
| Zdr-6 | Czech | CS22589 | Jirina<br>Relichová | 5/31/2000 | Czech<br>Republic | 49.3853 | 16.2544 | Zdr |

**Supplementary Table 2.** List of 96 *P.rapae* individuals used in Pieris GWAS.

| **Sample ID** | **Location** | **# Reads** | **SNPs per sample** |
| --- | --- | --- | --- |
| SN1-1 | University of Chicago,IL | 16,835,642 | 3,831,390 |
| SN1-2 | University of Chicago,IL | 16,690,476 | 3,499,432 |
| SN1-3 | University of Chicago,IL | 26,957,598 | 4,262,071 |
| SN1-4 | University of Chicago,IL | 18,250,528 | 3,792,942 |
| SN1-5 | Schaumburg,IL | 28,371,122 | 3,778,816 |
| SN1-6 | University of Chicago,IL | 25,206,968 | 4,120,740 |
| SN1-7 | University of Chicago,IL | 23,495,440 | 4,199,475 |
| SN1-8 | University of Chicago,IL | 28,485,402 | 4,328,932 |
| SN1-9 | University of Chicago,IL | 15,116,196 | 3,686,801 |
| SN1-10 | University of Chicago,IL | 19,030,478 | 4,565,351 |
| SN1-11 | University of Chicago,IL | 27,052,932 | 5,358,852 |
| SN1-12 | University of Chicago,IL | 14,893,910 | 2,928,379 |
| SN1-13 | University of Chicago,IL | 18,059,212 | 3,828,798 |
| SN1-14 | University of Chicago,IL | 22,820,320 | 3,983,847 |
| SN1-15 | University of Chicago,IL | 36,021,418 | 4,356,063 |
| SN1-16 | University of Chicago,IL | 12,271,404 | 3,419,565 |
| SN1-17 | University of Chicago,IL | 26,675,546 | 4,189,629 |
| SN1-18 | University of Chicago,IL | 22,361,262 | 3,926,289 |
| SN1-19 | University of Chicago,IL | 18,815,332 | 3,919,399 |
| SN1-20 | University of Chicago,IL | 21,906,464 | 4,663,772 |
| SN1-21 | University of Chicago,IL | 18,279,550 | 3,771,071 |
| SN1-22 | University of Chicago,IL | 11,794,858 | 4,002,105 |
| SN1-23 | University of Chicago,IL | 18,078,476 | 3,817,351 |
| SN1-24 | University of Chicago,IL | 26,813,358 | 4,215,037 |
| SN2-1 | University of Chicago,IL | 35,502,330 | 4,404,469 |
| SN2-2 | University of Chicago,IL | 26,399,338 | 4,418,141 |
| SN2-3 | University of Chicago,IL | 12,114,480 | 3,369,810 |
| SN2-4 | University of Chicago,IL | 30,325,014 | 4,340,905 |
| SN2-5 | University of Chicago,IL | 28,937,674 | 4,257,918 |
| SN2-6 | University of Chicago,IL | 24,441,562 | 4,098,097 |
| SN2-7 | University of Chicago,IL | 14,409,634 | 3,298,740 |
| SN2-8 | University of Chicago,IL | 17,851,616 | 3,686,303 |
| SN2-9 | University of Chicago,IL | 29,575,908 | 4,242,049 |
| SN2-10 | University of Chicago,IL | 25,206,240 | 4,170,416 |
| SN2-11 | University of Chicago,IL | 12,842,508 | 3,354,444 |
| SN2-12 | University of Chicago,IL | 28,153,206 | 4,421,812 |
| SN2-13 | University of Chicago,IL | 16,721,526 | 3,738,070 |
| SN2-14 | University of Chicago,IL | 25,174,770 | 4,145,893 |
| SN2-15 | University of Chicago,IL | 23,191,436 | 3,917,609 |
| SN2-16 | University of Chicago,IL | 30,018,634 | 4,213,145 |
| SN2-17 | University of Chicago,IL | 23,429,212 | 4,058,390 |
| SN2-18 | University of Chicago,IL | 31,742,776 | 4,156,959 |
| SN2-19 | University of Chicago,IL | 24,448,420 | 4,052,921 |
| SN2-20 | University of Chicago,IL | 18,959,980 | 3,722,312 |
| SN2-21 | University of Chicago,IL | 39,691,768 | 4,830,783 |
| SN2-22 | University of Chicago,IL | 24,922,346 | 4,264,440 |
| SN2-23 | University of Chicago,IL | 39,987,340 | 4,401,123 |
| SN2-24 | University of Chicago,IL | 34,115,436 | 4,447,627 |
| SN2-25 | University of Chicago,IL | 16,194,606 | 3,468,528 |
| SN2-26 | University of Chicago,IL | 17,365,278 | 3,814,271 |
| SN2-27 | University of Chicago,IL | 15,122,652 | 4,306,601 |
| SN2-28 | Schaumburg,IL | 31,841,314 | 4,374,753 |
| SN2-29 | Schaumburg,IL | 17,495,634 | 3,694,208 |
| SN2-30 | Schaumburg,IL | 14,902,966 | 3,780,863 |
| SN2-31 | Schaumburg,IL | 15,245,638 | 3,839,257 |
| SN2-32 | University of Chicago,IL | 45,800,742 | 4,341,145 |
| SN2-33 | Schaumburg,IL | 24,533,058 | 4,160,031 |
| SN2-34 | Schaumburg,IL | 29,684,602 | 4,194,515 |
| SN2-35 | Schaumburg,IL | 27,851,996 | 4,270,839 |
| SN2-36 | Schaumburg,IL | 22,625,152 | 4,080,784 |
| SN2-37 | Schaumburg,IL | 14,306,364 | 3,057,695 |
| SN2-38 | Schaumburg,IL | 18,849,984 | 3,790,211 |
| SN2-39 | Schaumburg,IL | 23,002,558 | 3,844,742 |
| SN2-40 | Schaumburg,IL | 15,183,264 | 3,780,936 |
| SN3-1 | Schaumburg,IL | 15,620,068 | 3,785,333 |
| SN3-2 | Schaumburg,IL | 29,512,116 | 4,169,538 |
| SN3-3 | Schaumburg,IL | 21,037,624 | 3,917,810 |
| SN3-4 | University of Chicago,IL | 24,676,850 | 4,088,482 |
| SN3-5 | Schaumburg,IL | 14,681,648 | 3,434,508 |
| SN3-6 | Chicago downtown,IL | 21,271,010 | 4,081,587 |
| SN3-7 | Schaumburg,IL | 14,866,824 | 3,781,025 |
| SN3-8 | University of Chicago,IL | 17,898,040 | 3,798,314 |
| SN3-9 | Schaumburg,IL | 17,474,592 | 3,845,304 |
| SN3-10 | University of Chicago,IL | 21,455,300 | 4,018,951 |
| SN3-11 | Carolina Biological<br>Supply,NC | 18,661,246 | 3,221,765 |
| SN3-12 | Chicago downtown,IL | 25,666,124 | 4,148,584 |
| SN3-13 | University of Chicago,IL | 19,371,456 | 3,879,331 |
| SN3-14 | University of Chicago,IL | 38,173,414 | 4,341,148 |
| SN3-15 | Chicago downtown,IL | 17,197,960 | 3,831,755 |
| SN3-16 | University of Chicago,IL | 19,148,234 | 3,923,953 |
| SN3-17 | Langdon,ND | 10,667,446 | 3,327,261 |
| SN3-18 | Langdon,ND | 15,693,070 | 3,689,625 |
| SN3-19 | Langdon,ND | 20,140,606 | 3,912,035 |
| SN3-20 | Langdon,ND | 34,623,120 | 4,438,222 |
| SN3-21 | Langdon,ND | 23,224,660 | 4,108,532 |
| SN3-22 | Langdon,ND | 20,277,212 | 3,997,092 |
| SN3-23 | Langdon,ND | 21,832,814 | 3,815,381 |
| SN3-24 | Langdon,ND | 17,669,434 | 3,513,475 |
| SN3-25 | Langdon,ND | 10,645,710 | 3,397,642 |
| SN3-26 | Langdon,ND | 16,729,946 | 3,739,124 |
| SN3-27 | Langdon,ND | 15,135,064 | 3,651,243 |
| SN3-28 | Langdon,ND | 28,124,188 | 4,454,347 |
| SN3-29 | Langdon,ND | 34,790,408 | 4,421,381 |
| SN3-30 | Langdon,ND | 18,450,192 | 3,960,058 |
| SN3-31 | Langdon,ND | 11,127,958 | 3,287,501 |
| SN3-32 | Langdon,ND | 15,590,884 | 3,720,455 |

**Supplementary Table 3.** List of various time points for RNAseq sample collection.

|  |  |  |
| --- | --- | --- |
| Time points | Host Plants | Butterflies |
| 72 h after oviposition | leaves with eggs | eggs deposited on leaves |
|  | control leaves without eggs | eggs deposited on wax paper |
| 24 h of larval feeding, age of larvae ~48h | leaves after 24 h of larval feeding from plants previously exposed to eggs | larvae after 24 h of feeding on leaves from plants exposed to eggs |
|  | leaves after 24 h of larval feeding from plants not previously exposed to eggs | larvae after 24 h of feeding on leaves from plants not exposed to eggs |
|  | control leaves never exposed to eggs or larvae | larvae after 24 h of feeding on artificial diet |
| 24 h after mechanical wounding | leaves 24 h after mechanical wounding | n/a |

**Supplementary Table 4.**
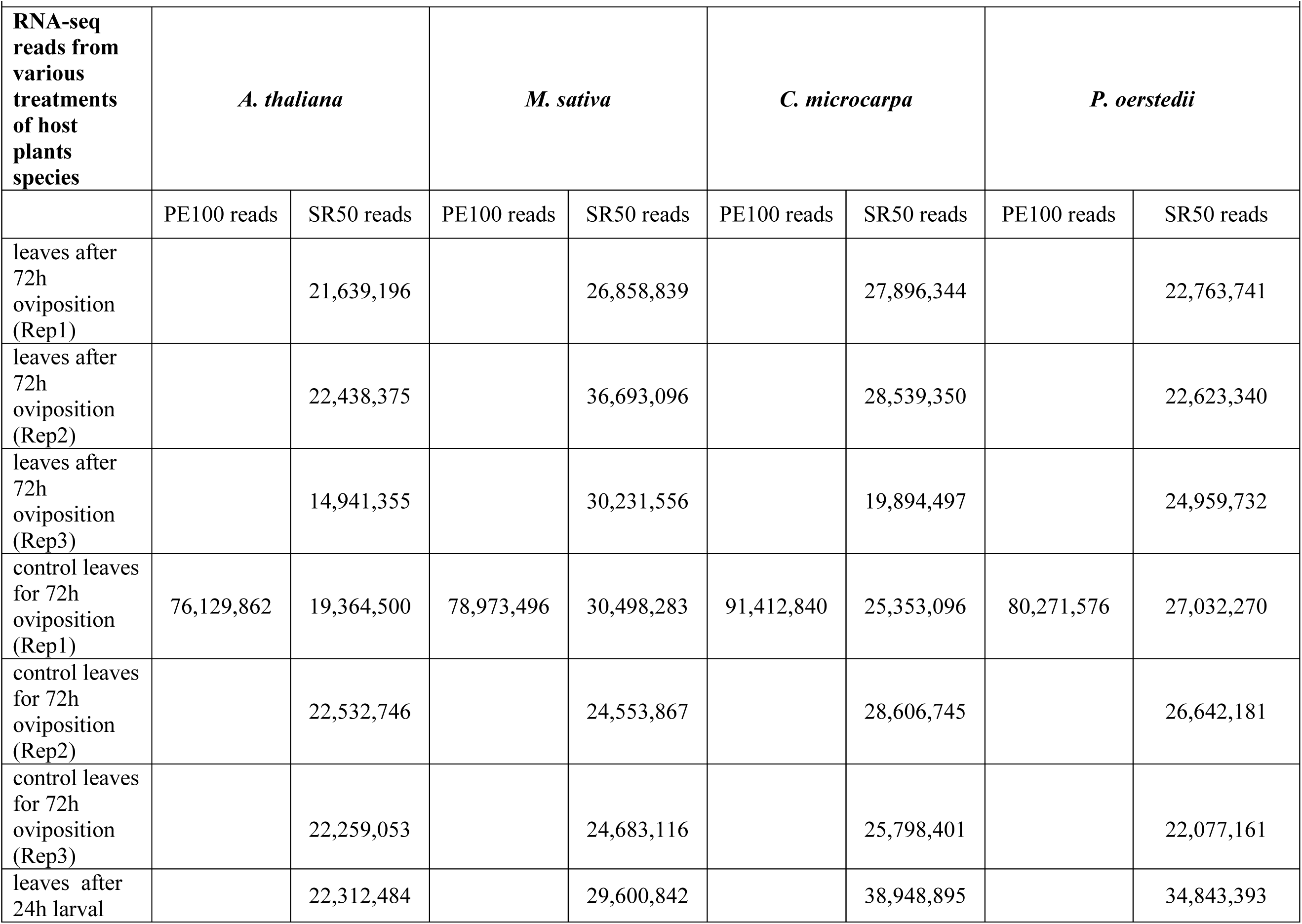
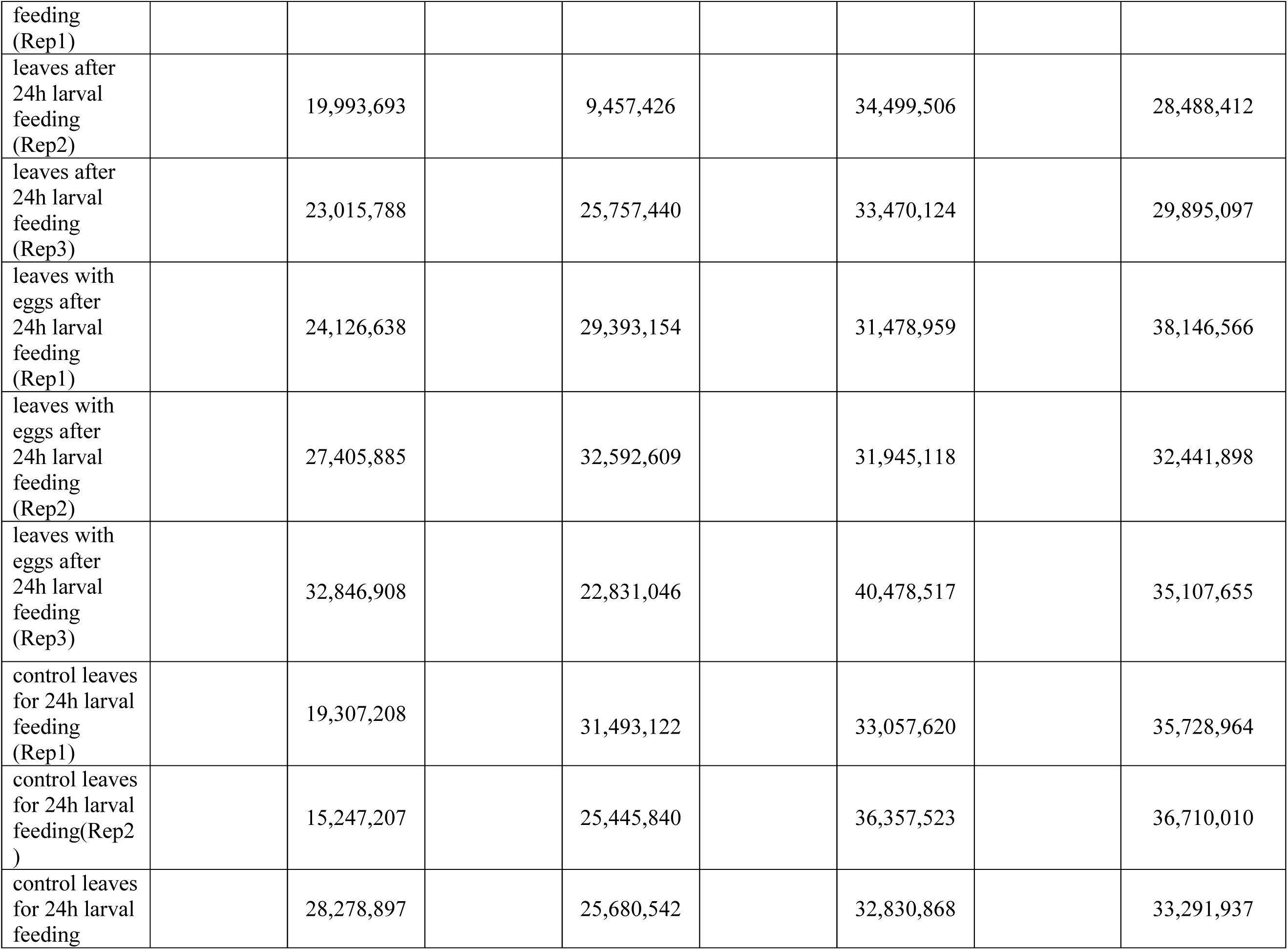
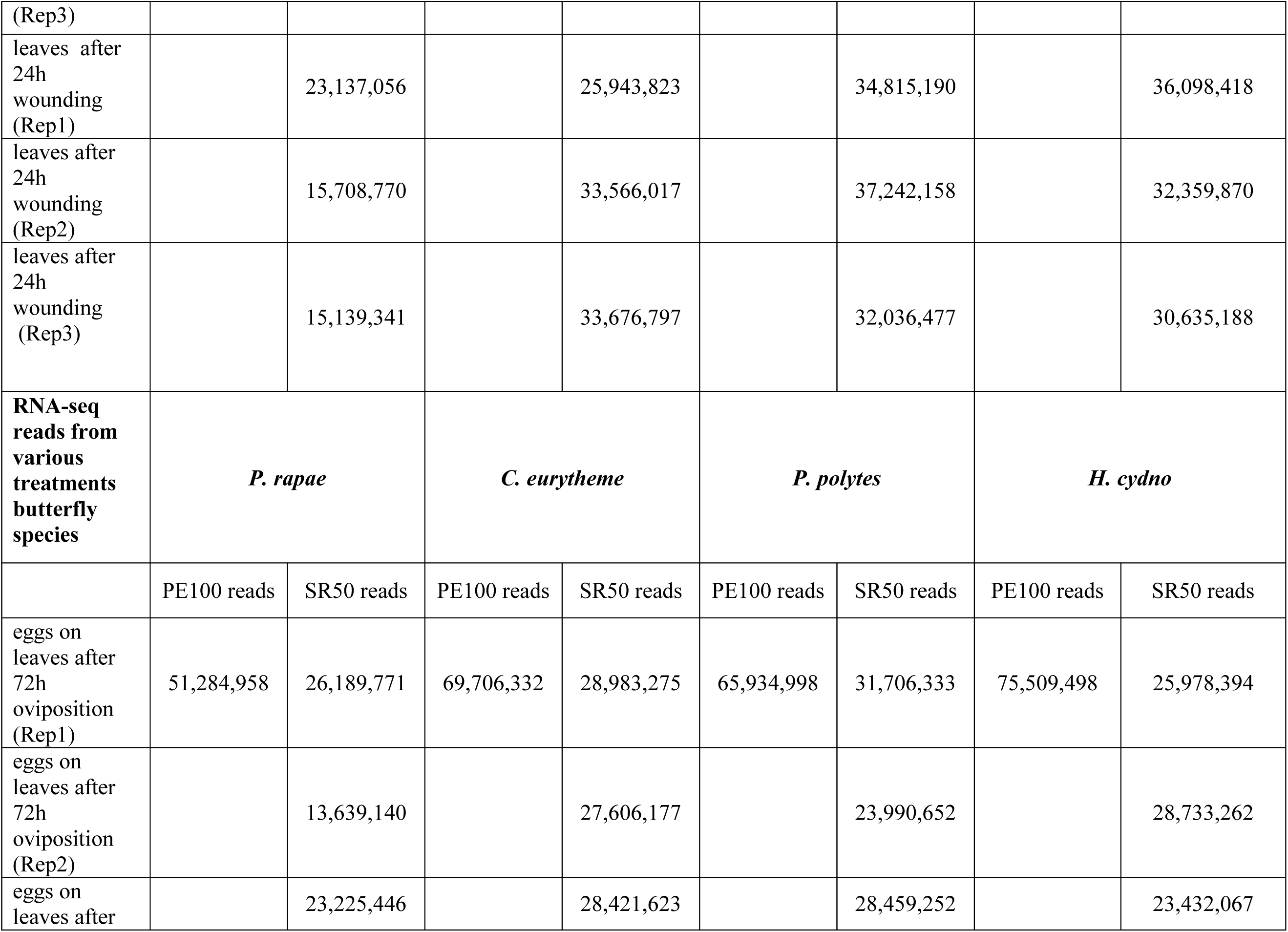
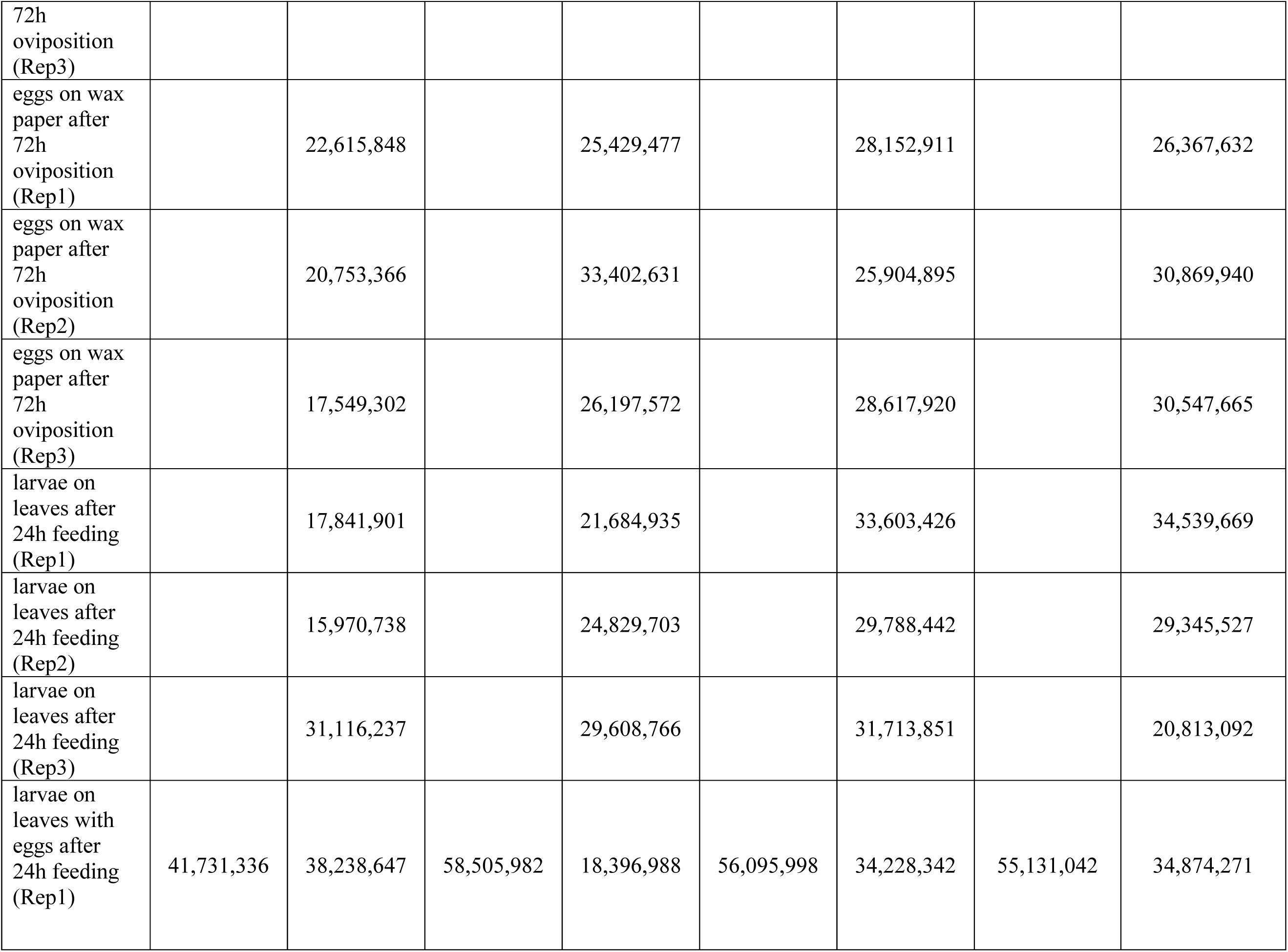
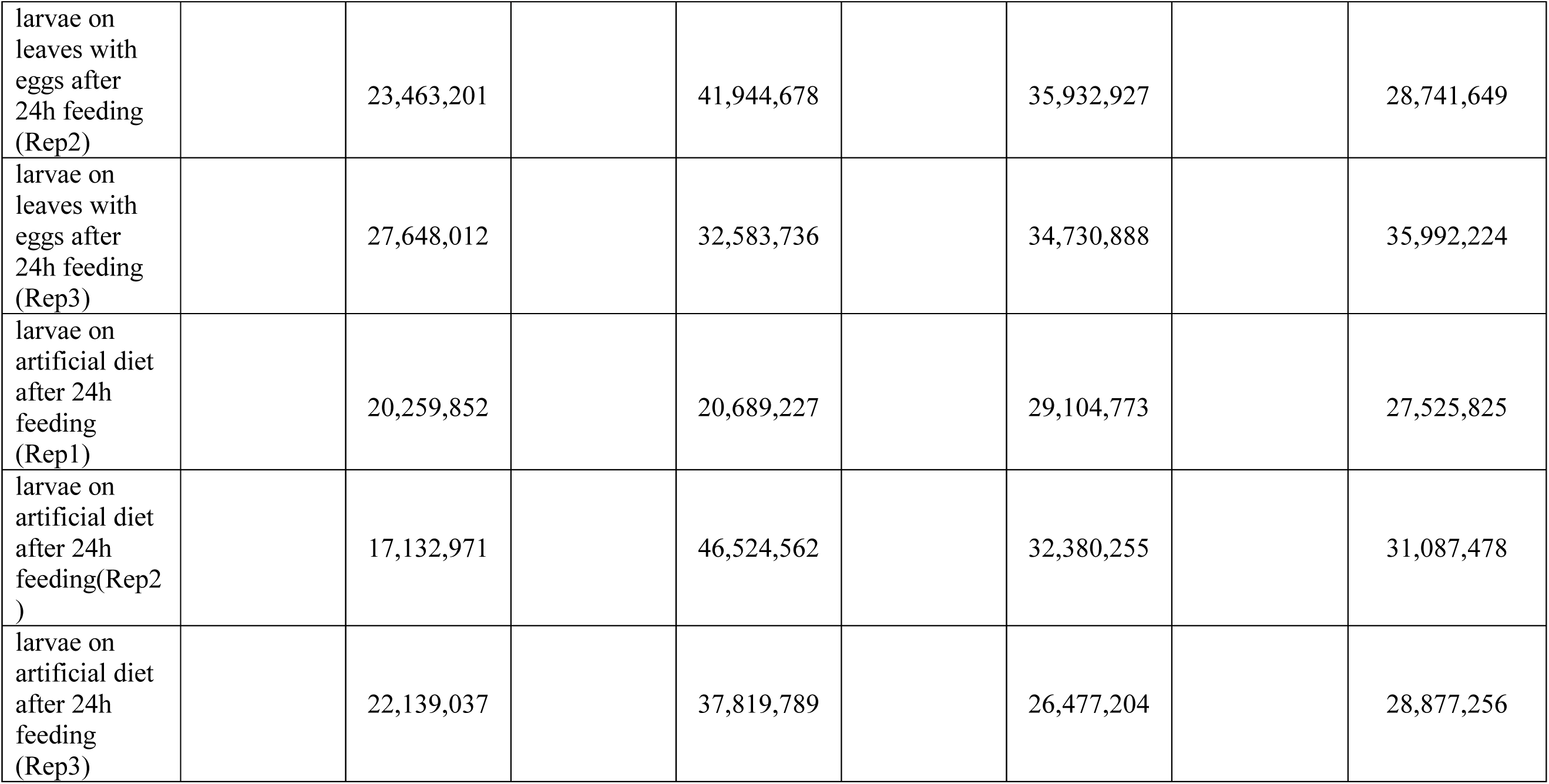
RNAseq reads from various treatments of host plants and butterfly species.

**Supplementary Table 5.**
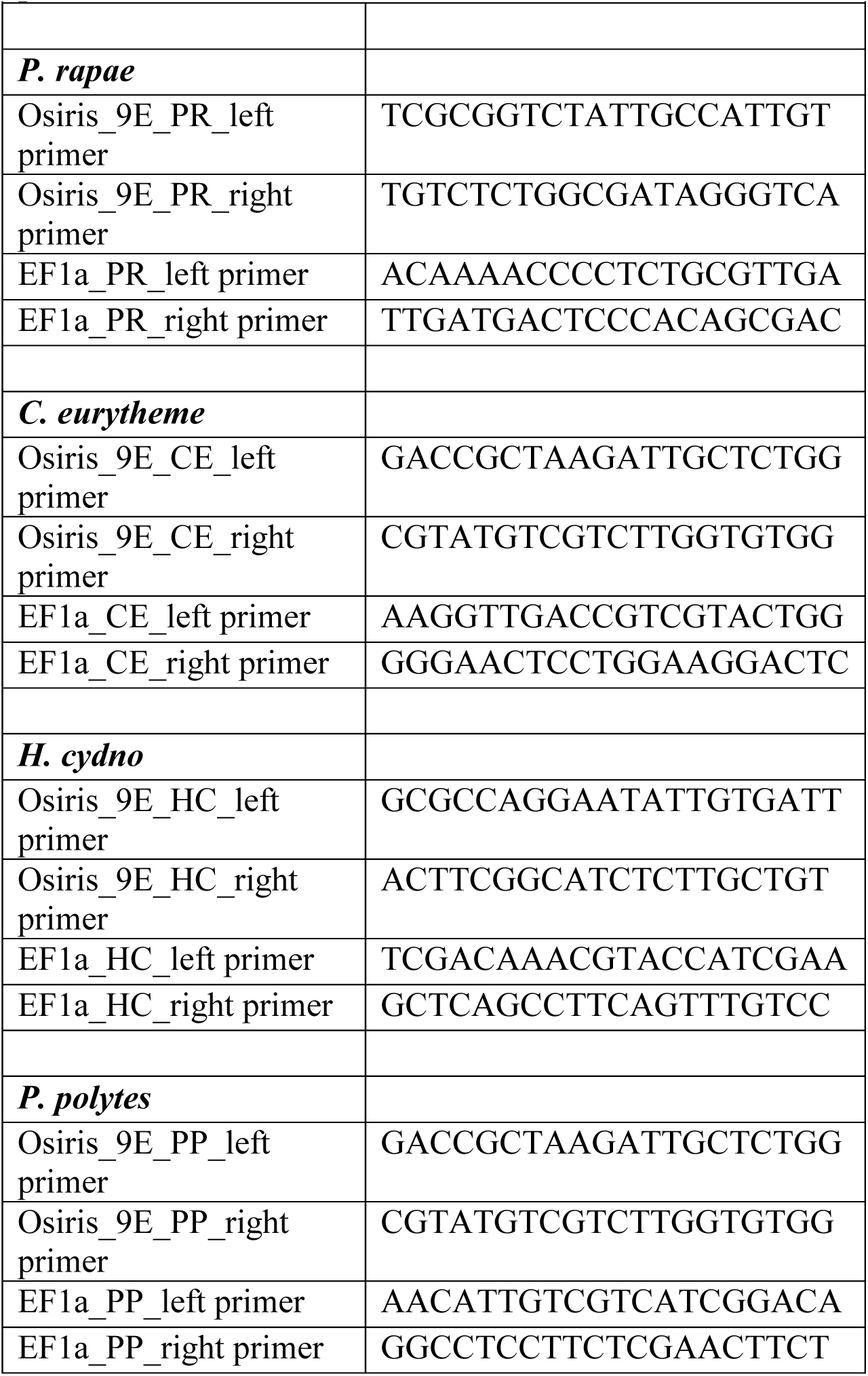
Primer sequences for cDNA synthesis for *Osiris 9E* spatial and temporal expression patterns.

## References

1. Futuyma, D.J. Some current approaches to the evolution of plant-herbivore interactions. Plant Species Biology 15, 1-9 (2000).

2. Futuyma, D.J. & Agrawal, A.A. Macroevolution and the biological diversity of plants and herbivores. Proc. Natl. Acad. Sci. USA 106, 18054-18061 (2009).

3. Ehrlich, P.R. & Raven, P.H. Butterflies and plants: A study in coevolution. Evolution 18, 586-608 (1964).

4. Howe, G.A. & Jander, G. Plant immunity to insect herbivores. Annu. Rev. Plant. Biol. 59, 41-66 (2008).

5. Berenbaum, M.R. Postgenomic chemical ecology: from genetic code to ecological interactions. J. Chem. Ecol. 28, 873-896 (2002).

6. Li, W., Schuler, M.A. & Berenbaum, M.R. Diversification of furanocoumarin-metabolizing cytochrome P450 monooxygenases in two papilionids: Specificity and substrate encounter rate. Proc. Natl. Acad. Sci. USA 100 Suppl 2, 14593-14598 (2003).

7. Wittstock, U. et al. Successful herbivore attack due to metabolic diversion of a plant chemical defense. Proc. Natl. Acad. Sci. USA 101, 4859-4864 (2004).

8. Petschenka, G. et al. Stepwise evolution of resistance to toxic cardenolides via genetic substitutions in the Na+/K+ -ATPase of milkweed butterflies (Lepidoptera: Danaini). Evolution 67, 2753-2761 (2013).

9. Groen, S.C. & Whiteman, N.K. Using *Drosophila* to study the evolution of herbivory and diet specialization. Curr. Opin. Insect Sci. 14, 66-72 (2016).

10. Schuman, M.C. & Baldwin, I.T. The layers of plant responses to insect herbivores. Annu. Rev. Entomol. 61, 373-394 (2016).

11. Mikkelsen, M.D., Hansen, C.H., Wittstock, U. & Halkier, B.A. Cytochrome P450 CYP79B2 from *Arabidopsis* catalyzes the conversion of tryptophan to indole-3- acetaldoxime, a precursor of indole glucosinolates and indole-3-acetic acid. J. Biol. Chem. 275, 33712-33717 (2000).

12. Kim, J.I., Dolan, W.L., Anderson, N.A. & Chapple, C. Indole glucosinolate biosynthesis limits phenylpropanoid accumulation in *Arabidopsis thaliana*. Plant Cell 27, 1529-1546 (2015).

13. Rajniak, J., Barco, B., Clay, N.K. & Sattely, E.S. A new cyanogenic metabolite in *Arabidopsis* required for inducible pathogen defence. Nature 525, 376-379 (2015).

14. Hopkins, R.J., van Dam, N.M. & van Loon, J.J. Role of glucosinolates in insect-plant relationships and multitrophic interactions. Annu. Rev. Entomol. 54, 57-83 (2009).

15. Wheat, C.W. et al. The genetic basis of a plant-insect coevolutionary key innovation. Proc. Natl. Acad. Sci. USA 104, 20427-20431 (2007).

16. Edger, P.P. et al. The butterfly plant arms-race escalated by gene and genome duplications. Proc. Natl. Acad. Sci. USA 112, 8362-8366 (2015).

17. Li, J., Zhao, J., Rose, A.B., Schmidt, R. & Last, R.L. *Arabidopsis* phosphoribosylanthranilate isomerase: molecular genetic analysis of triplicate tryptophan pathway genes. Plant Cell 7, 447-461 (1995).

18. Bartels, S. et al. The family of Peps and their precursors in *Arabidopsis*: differential expression and localization but similar induction of pattern-triggered immune responses. J. Exp. Bot. 64, 5309-5321 (2013).

19. Brachi, B. et al. Coselected genes determine adaptive variation in herbivore resistance throughout the native range of *Arabidopsis thaliana*. Proc. Natl. Acad. Sci. USA 112, 4032-4037 (2014).

20. Maeki, K. & Remignton, C. Studies of the chromosomes of North American Rhopalocera. J. Lep. Soc. 14, 37-56 (1960).

21. Noctor, G., Queval, G., Mhamdi, A., Chaouch, S. & Foyer, C.H. Glutathione. Arabidopsis Book 9, e0142 (2011).

22. Nelson, R.E. et al. Peroxidasin: A novel enzyme-matrix protein of *Drosophila* development. EMBO J. 13, 3438-3447 (1994).

23. Soudi, M., Zamocky, M., Jakopitsch, C., Furtmüller, P.G. & Obinger, C. Molecular evolution, structure, and function of peroxidasins. Chem. Biodivers 9, 1776-1793 (2012).

24. Hilker, M. & Fatouros, N.E. Plant responses to insect egg deposition. Annu. Rev. Entomol. 60, 493-515 (2015).

25. Jones, J.D. & Dangl, J.L. The plant immune system. Nature 444, 323-329 (2006).

26. Little, D., Gouhier-Darimont, C., Bruessow, F. & Reymond, P. Oviposition by pierid butterflies triggers defense responses in *Arabidopsis*. Plant Physiol. 143, 784-800 (2007).

27. Gouhier-Darimont, C., Schmiesing, A., Bonnet, C., Lassueur, S. & Reymond, P. Signalling of *Arabidopsis thaliana* response to *Pieris brassicae* eggs shares similarities with PAMP-triggered immunity. J. Exp. Bot. 64, 665-674 (2013).

28. Schmiesing, A., Emonet, A., Gouhier-Darimont, C. & Reymond, P. *Arabidopsis* MYC transcription factors are the target of hormonal salicylic acid/jasmonic acid cross talk in response to *Pieris brassicae* egg extract. Plant Physiol. 170, 2432-2443 (2016).

29. Zou, X. et al. Glutathione s-transferase SlGSTE1 in *Spodoptera litura* may be associated with feeding adaptation of host plants. Insect Biochem. Mol. Biol. 70, 32-43 (2016).

30. Yu, Q. et al. The transcriptome response of *Heliconius melpomene* larvae to a novel host plant. Mol. Ecol. 25, 4850-4865 (2016).

31. Schweizer, F. et al. *Arabidopsis* glucosinolates trigger a contrasting transcriptomic response in a generalist and a specialist herbivore. Insect Biochem. Mol. Biol. 85, 21-31 (2017).

32. Gloss, A. D. et al. Evolution in an ancient detoxification pathway is coupled with a transition to herbivory in the Drosophilidae. Mol. Biol. Evol. 31, 2441-2456 (2014).

33. Agrios, G. Plant Pathology, 952 (Academic Press; 5 edition, 2005).

34. Tsuda, K. & Somssich, I.E. Transcriptional networks in plant immunity. New Phytol. 206, 932-947 (2015).

35. Roitberg, B.D. & Isman, M.B. Insect Chemical Ecology, 360 (Springer US, 1992).

36. Gerardo, N.M. et al. Immunity and other defenses in pea aphids, *Acyrthosiphon pisum*. Genome Biol. 11, R21 (2010).

37. Shah, N., Dorer, D.R., Moriyama, E.N. & Christensen, A.C. Evolution of a large, conserved, and syntenic gene family in insects. G3 2, 313–319 (2012).

38. Dorer, D.R., Rudnick, J.A., Moriyama, E.N. & Christensen, A.C. A family of genes clustered at the triplo-lethal locus of *Drosophila melanogaster* has an unusual evolutionary history and significant synteny with *Anopheles gambiae*. Genetics 165, 613–621 (2003).

39. Hungate, E.A. et al. A locus in *Drosophila sechellia* affecting tolerance of a host plant toxin. Genetics 195, 1063–1075 (2013).

40. Andrade López, J.M. et al. Genetic basis of octanoic acid resistance in *Drosophila sechellia*: functional analysis of a fine-mapped region. Mol. Ecol. 26, 1148-1160 (2017).

41. Yassin, A. et al. Recurrent specialization on a toxic fruit in an island *Drosophila* population. Proc. Natl. Acad. Sci. USA 113, 4771–4776 (2016).

42. Vanderauwera, S. et al. Genome-wide analysis of hydrogen peroxide-regulated gene expression in *Arabidopsis* reveals a high light-induced transcriptional cluster involved in anthocyanin biosynthesis. Plant Physiol. 139, 806-821 (2005).

43. Queval, G. & Noctor, G. A plate reader method for the measurement of NAD, NADP, glutathione, and ascorbate in tissue extracts: Application to redox profiling during *Arabidopsis* rosette development. Anal. Biochem. 363, 58-69 (2007).

44. Queval, G. et al. H2O2-activated up-regulation of glutathione in *Arabidopsis* involves induction of genes encoding enzymes involved in cysteine synthesis in the chloroplast. Mol. Plant 2, 344-356 (2009).

45. Chaouch, S. & Noctor, G. Myo-inositol abolishes salicylic acid-dependent cell death and pathogen defence responses triggered by peroxisomal hydrogen peroxide. New Phytol. 188, 711-718 (2010).

46. Parisy, V. et al. Identification of PAD2 as a gamma-glutamylcysteine synthetase highlights the importance of glutathione in disease resistance of *Arabidopsis*. Plant J. 49, 159-172 (2007).

47. Schlaeppi, K., Bodenhausen, N., Buchala, A., Mauch, F. & Reymond, P. The glutathione-deficient mutant pad2-1 accumulates lower amounts of glucosinolates and is more susceptible to the insect herbivore *Spodoptera littoralis*. Plant J. 55, 774-786 (2008).

48. Loudet, O. et al. Natural variation for sulfate content in *Arabidopsis thaliana* is highly controlled by APR2. Nat. Genet. 39, 896-900 (2007).

49. Whiteman, N.K. et al. Genes involved in the evolution of herbivory by a leaf-mining, Drosophilid fly. Genome Biol. Evol. 4, 900-916 (2012).

50. Green, M. & Sambrook, J. Molecular Cloning: A Laboratory Manual (Fourth Edition), 2028 (Cold Spring Harbor Laboratory Press, 2012).

51. Putnam, N.H. et al. Chromosome-scale shotgun assembly using an in vitro method for long-range linkage. Genome Res. 26, 342-350 (2016).

52. Marçais, G. & Kingsford, C. A fast, lock-free approach for efficient parallel counting of occurrences of k-mers. Bioinformatics 27, 764-770 (2011).

53. Leggett, R.M., Clavijo, B.J., Clissold, L., Clark, M.D. & Caccamo, M. NextClip: an analysis and read preparation tool for Nextera Long Mate Pair libraries. Bioinformatics 30, 566-568 (2014).

54. Gnerre, S. et al. High-quality draft assemblies of mammalian genomes from massively parallel sequence data. Proc. Natl. Acad. Sci. USA 108, 1513-1518 (2011).

55. Parra, G., Bradnam, K. & Korf, I. CEGMA: a pipeline to accurately annotate core genes in eukaryotic genomes. Bioinformatics 23, 1061-1067 (2007).

56. Boetzer, M., Henkel, C.V., Jansen, H.J., Butler, D. & Pirovano, W. Scaffolding pre-assembled contigs using SSPACE. Bioinformatics 27, 578-579 (2011).

57. Kiełbasa, S.M., Wan, R., Sato, K., Horton, P. & Frith, M.C. Adaptive seeds tame genomic sequence comparison. Genome Res. 21, 487-493 (2011).

58. Hill, J. et al. A butterfly genome reveals extensive chromosomal reshuffling. In preparation.

59. Magrane, M. & Consortium, U. UniProt Knowledgebase: a hub of integrated protein data. Database (Oxford) 2011, bar009 (2011).

60. Meslin, C. et al. Digestive organ in the female reproductive tract borrows genes from multiple organ systems to adopt critical functions. Mol. Biol. Evol. 32, 1567-1580 (2015).

61. Trapnell, C. et al. Differential gene and transcript expression analysis of RNA-seq experiments with TopHat and Cufflinks. Nat. Protoc. 7, 562-578 (2012).

62. Kim, D. et al. TopHat2: accurate alignment of transcriptomes in the presence of insertions, deletions and gene fusions. Genome Biol. 14, R36 (2013).

63. Trapnell, C. et al. Transcript assembly and quantification by RNA-Seq reveals unannotated transcripts and isoform switching during cell differentiation. Nat. Biotechnol. 28, 511-515 (2010).

64. Holt, C. & Yandell, M. MAKER2: an annotation pipeline and genome-database management tool for second-generation genome projects. BMC Bioinformatics 12, 491 (2011).

65. Stanke, M., Diekhans, M., Baertsch, R. & Haussler, D. Using native and syntenically mapped cDNA alignments to improve de novo gene finding. Bioinformatics 24, 637-644 (2008).

66. Horton, M.W. et al. Genome-wide patterns of genetic variation in worldwide *Arabidopsis thaliana* accessions from the RegMap panel. Nat. Genet. 44, 212-216 (2012).

67. Zhou, X. & Stephens, M. Efficient multivariate linear mixed model algorithms for genome-wide association studies. Nat. Methods 11, 407-409 (2014).

68. Bolger, A.M., Lohse, M. & Usadel, B. Trimmomatic: a flexible trimmer for Illumina sequence data. Bioinformatics 30, 2114-2120 (2014).

69. DePristo, M.A. et al. A framework for variation discovery and genotyping using next-generation DNA sequencing data. Nat. Genet. 43, 491-498 (2011).

70. Van der Auwera, G.A. et al. From FastQ data to high confidence variant calls: the Genome Analysis Toolkit best practices pipeline. Curr. Protoc. Bioinformatics 43, 11.10.1-11.10.33 (2013).

71. Gao, X. Multiple testing corrections for imputed SNPs. Genet. Epidemiol. 35, 154-158 (2011).

72. Dobin, A. et al. STAR: ultrafast universal RNA-seq aligner. Bioinformatics 29, 15-21 (2013).

73. Haas, B.J. et al. De novo transcript sequence reconstruction from RNA-seq using the Trinity platform for reference generation and analysis. Nat. Protoc. 8, 1494-1512 (2013).

74. Conesa, A. et al. Blast2GO: a universal tool for annotation, visualization and analysis in functional genomics research. Bioinformatics 21, 3674-3676 (2005).

75. Lechner, M. et al. Proteinortho: detection of (co-)orthologs in large-scale analysis. BMC Bioinformatics 12, 124 (2011).

76. Lechner, M. et al. Orthology detection combining clustering and synteny for very large datasets. PLoS One 9, e105015 (2014).

77. Katoh, K. & Toh, H. Recent developments in the MAFFT multiple sequence alignment program. Brief. Bioinform. 9, 286–298 (2008).

78. Price, M.N. et al. FastTree: Computing large minimum-evolution trees with profiles instead of a distance matrix. Mol. Biol. Evol. 26, 1641-1650 (2009).

79. Heikkilä, M., Kaila, L., Mutanen, M., Peña, C. & Wahlberg, N. Cretaceous origin and repeated tertiary diversification of the redefined butterflies. Proc. R. Soc. Lond. B. 279, 1093-1099 (2012).

